# FIT2 is a lipid phosphate phosphatase crucial for endoplasmic reticulum homeostasis

**DOI:** 10.1101/291765

**Authors:** Michel Becuwe, Laura M. Bond, Niklas Mejhert, Sebastian Boland, Shane D. Elliott, Marcelo Cicconet, Xinran N. Liu, Morven M. Graham, Tobias C. Walther, Robert V. Farese

## Abstract

The endoplasmic reticulum (ER) protein Fat-Induced Transcript 2 (FIT2) has emerged as a key factor in lipid droplet (LD) formation, although its molecular function is unknown. Highlighting its importance, FIT2 orthologs are essential in worms and mice, and FIT2 deficiency causes a deafness/dystonia syndrome in humans. Here we show that FIT2 is a lipid phosphate phosphatase (LPP) enzyme that is required for maintaining the normal structure of the ER. Recombinant FIT2 exhibits LPP activity *in vitro* and loss of this activity in cells leads to ER membrane morphological changes and ER stress. Defects in LD formation in FIT2 depletion appear to be secondary to membrane lipid abnormalities, possibly due to alterations in phospholipids required for coating forming LDs. Our findings uncover an enzymatic role for FIT2 in ER lipid metabolism that is crucial for ER membrane homeostasis.

## Introduction

The endoplasmic reticulum (ER) is a network of membrane tubules and sheets that span the entire cell and contain ~50% of cell membranes. Key functions of the ER include the synthesis of most secreted and membrane proteins and of each of the major classes of lipids (e.g., phospholipids, sterols, sphingolipids) in the cell. Proteins, such as reticulons (De Craene et al., 2006; Voeltz et al., 2006) and atlastins (Hu et al., 2009; Orso et al., 2009), are key players in shaping and fusing ER tubules, respectively, and other proteins have been implicated in sheet formation (Shibata et al., 2010).

Lipid droplet (LD) formation is an important function of the ER (Choudhary et al., 2015; Jacquier et al., 2011; Kassan et al., 2013). These cytosolic organelles serve as cellular reservoirs for neutral lipids that are used for membranes or generating metabolic energy (Walther and Farese, 2012). How LDs are formed in the ER is a fundamental and unsolved problem in cell biology. A current model posits that neutral lipids, such as triacylglycerols (TGs), are formed in the ER through the action of enzymes, such as the DGAT enzymes (Cases and Farese, 1998; Cases et al., 2001; Lardizabal et al., 2001). The neutral lipids are thought to diffuse in the ER bilayer and, when they reach a critical concentration, to nucleate a neutral-lipid lens within the bilayer. This lens grows and eventually a LD buds towards the cytosol. After reaching a specific size (typical initial LDs are 300–600 nm diameter), the forming LD may detach from the ER bilayer (Walther et al., 2016).

The phospholipid composition of the ER is a key parameter in controlling LD budding (Ben M’barek et al., 2017; Chorlay and Thiam, 2018). In addition, evidence suggests LD formation is governed and organized by specific proteins (Walther et al., 2016). Among such proteins, the Fat-Inducible Transcript (FIT) proteins, FIT1 and FIT2, have emerged as key players (Kadereit et al., 2008). FIT1 is expressed predominantly in muscle cells in humans, whereas FIT2 is expressed broadly and at relatively high levels in adipose tissue (Kadereit et al., 2008). FIT2 was originally identified as a transcript that was induced after treatment with PPARα agonist in mice (Kadereit et al., 2008). Consistent with its expression pattern, FIT2 is important in LD formation and lipid storage (Choudhary et al., 2015; Kadereit et al., 2008) in all model systems investigated, including yeast, fish, worms and mice (Choudhary et al., 2015; Kadereit et al., 2008). In *Caenorhabditis elegans*, deletion of the FIT2 orthologue is not compatible with life (Choudhary et al., 2015). The global postnatal knockout of FIT2 in mice also results in lethality due to catastrophic effects on the small intestine (Goh et al., 2015). The selective knockout of murine FIT2 in adipose tissue leads to a progressive lipodystrophy (Miranda et al., 2014). Homozygous FIT2 deficiency in humans was recently reported to cause deafness-dystonia (Zazo Seco et al., 2017). These results indicate that FIT2 has a crucial role in cellular lipid metabolism. The molecular function of FIT2 is a mystery. Human FIT2 protein binds TG and diacylglycerol (Gross et al., 2011), but it is not strictly required for TG synthesis. Yeast orthologs of FIT2, *SCS3* and *YFT2*, have synthetic-genetic interactions suggestive of defects in phospholipid metabolism (Moir et al., 2012). Supporting this, *scs3*Δ cells are auxotrophic for inositol, a condition exacerbated by addition of choline, both key metabolites in phospholipid synthesis (Hosaka et al., 1994). Intriguingly, Scs3 and Yft2 have been implicated as crucial determinants for the directionality of LD budding (Choudhary et al., 2015). These previous observations led to a model in which FIT2 is predominantly a structural protein, with neutral lipid binding, that may be involved in the partitioning of lipids for LD formation and budding (Gross et al., 2011; Kadereit et al., 2008).

In the current study, we investigated FIT2 function in mammalian and yeast cells. Our studies uncover an enzymatic function for FIT2 as a lipid phosphate phosphatase (LPP) at the ER, where its activity is required to maintain ER homeostasis.

## Results

### FIT2 is required for normal LD formation in mammalian cells

Because FIT2 is required for normal LD formation and directionality of LD budding in yeast and for LD formation in murine fibroblasts (Choudhary et al., 2015; Kadereit et al., 2008), we tested whether FIT2 is involved in LD formation in human SUM159 cells. We first examined FIT2 localization during LD formation, at early and late time points, in cells expressing an N-terminal GFP-tagged version of FIT2. We found that FIT2 displayed a diffuse signal that co-localized with the ER marker signal sequence-BFP-KDEL, and this localization appeared unchanged during LD formation (Figure 1A).

**Figure 1:**
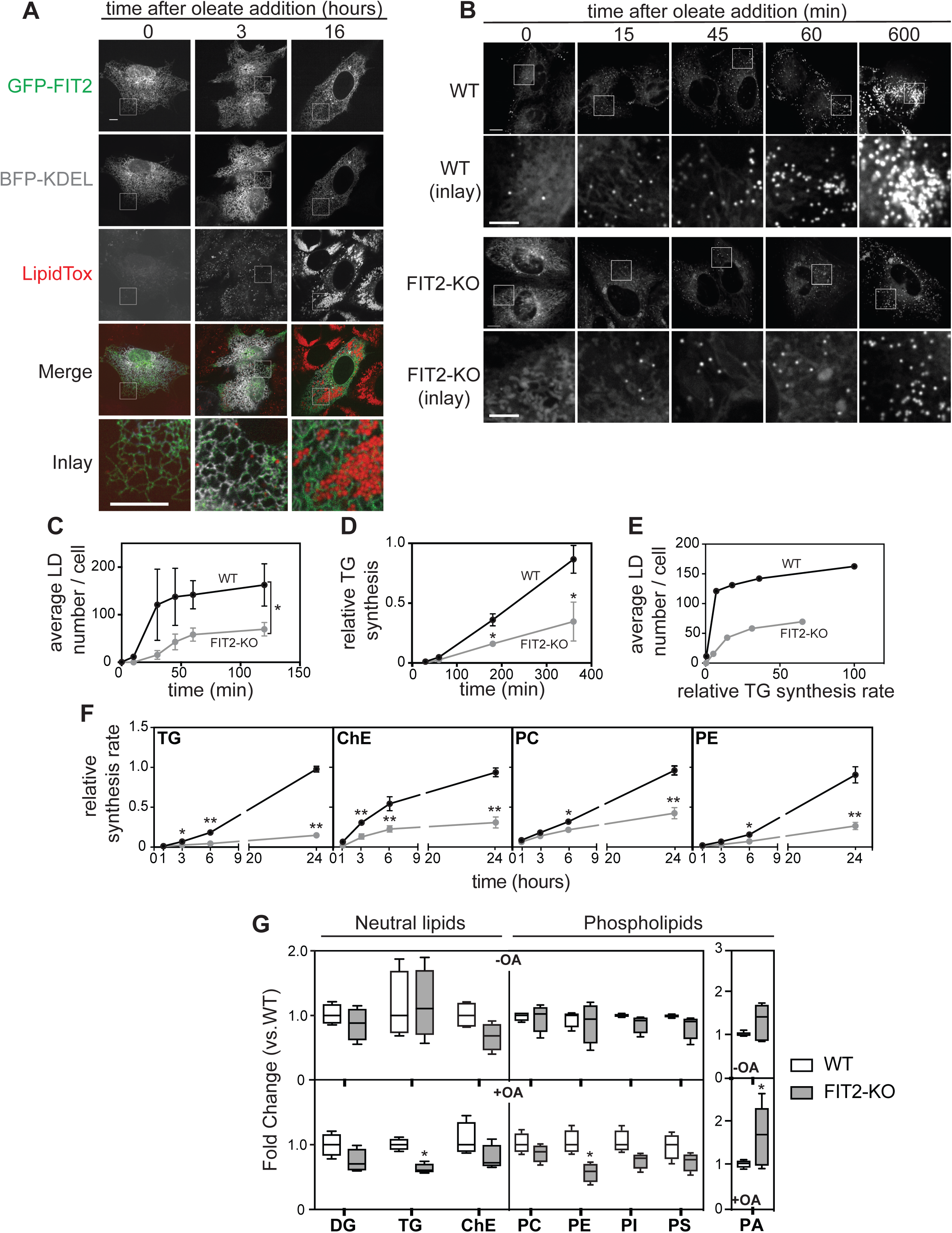
FIT2 is an ER protein required for normal LD formation and lipid synthesis in human cells. (**A**) FIT2 localizes diffusely throughout the ER. Wild-type SUM159 cells co-transfected with plasmids expressing the luminal ER marker ssBFP-KDEL and GFP-FIT2 were imaged before and after addition of oleic acid at the indicated time. LDs were stained with LipidTox. Bar, 10 µm. (**B**) FIT2 deletion results in fewer and smaller LDs formed in response to oleate treatment. Time course of LD formation in wild-type and FIT2-KO SUM159 cells. Cells were treated with oleic acid for the indicated times, and LDs were stained with BODIPY 493/503. Bar, 10 µm. (**C**) Quantification of LD formation in (B). n=4 cells. Values represent mean ± S.D (two-way ANOVA, \**p*<0.001). (**D**) Reduced triacylglycerol synthesis in FIT2-KO cells. Cells were pulse-labeled with [^14^C]-oleic acid, and incorporation into triacylglycerol (TG) was measured over time after oleate loading by TLC. Mean value of TG band intensity was plotted over time and quantified from three independent measurements ± S.D. Values were calculated relative to wild-type cells highest value at 360min (two-way ANOVA, Sidak Test, \**p*<0.0001). (**E**) FIT2-deficient cells form fewer LDs for similar amount of synthesized TG. Average LD number per cell measured in (E) was plotted against TG amount calculated in (D). (**F**) Reduced incorporation of [^14^C]-oleic acid during triacylglycerol (TG), cholesterol ester (ChE) phosphatidylcholine (PC) and phosphatidylethanolamine (PE) synthesis in FIT2-KO cells. Cells were pulse-labeled with [^14^C]-oleic acid and incorporation into the indicated lipids was measured over time during oleate loading by TLC. Mean value of band intensity was plotted over time and quantified from three independent measurements ± S.D. Values were calculated relative to 24 h time point value of wild-type cells (two-way ANOVA, Sidak Test, \**p*<0.05, \*\**p*<0.001). (**G**) Global analyses of wild-type and FIT2-KO cells lipidomes. Diacylglycerol (DG), triacylglycerol (TG), cholesterol ester (ChE), phosphatidylcholine (PC), phosphatidylethanolamine (PE), phosphatidylinositol (PI), and phosphatidylserine (PS). Phosphatidic acid (PA) levels were analyzed separately as describe in Methods. Values represent median ± S.D (two-way ANOVA, Sidak Test, \**p*<0.05).

To assess FIT2 function, we generated FIT2 knockout (FIT2-KO) lines in SUM-159 cells by CRISPR/Cas9-directed gene targeting. Specifically, we generated a cell line that has two identical alleles with a 17-base pair deletion of *FIT2* sequence at the junction of exon 1 and the first intron (designated clone1; Figure S1C). These cells lacked detectable FIT2 protein (Figure S1A). Incubation of these FIT2-KO cells in medium containing oleate revealed that they had fewer LDs at each time point, compared with control cells (Figure 1B-C).

To determine if the reduction in LD number was due to altered synthesis of TG or a defect in LD biogenesis, we measured TG synthesis by including a radioactive tracer ([^14^C]oleate) in the oleate-containing medium. We found that FIT2 deletion impaired TG synthesis, compared with wild-type cells (Figure 1D), similar to what was reported in mouse 3T3-L1 cells with FIT2 knock-down (Kadereit et al., 2008). Analysis of the numbers of LDs formed versus the amount of TG synthesized for each time point showed that FIT2 deficiency resulted in fewer LDs formed for any given level of newly synthesized TG (Figure 1E). Significantly, the incorporation of [^14^C]oleate into other lipid species, such as phosphatidylcholine (PC), phosphatidylethanolamine (PE) and cholesterol ester (ChE), was also reduced in FIT2-KO cells (Figure 1F). Indeed, global analyses of wild-type and FIT2-KO cell lipidomes 24 h after oleate addition confirmed that FIT2-KO cells have reduced levels of neutral lipids and phospholipids upon oleate treatment (Figure 1G). In contrast, FIT2-KO cells had ~50% more phosphatidic acid (PA) than wild-type cells. No observed differences in the lipidomes were found in wild-type or FIT2-KO cells in the basal states.

Mammalian cells synthesize TG in reactions catalyzed by either DGAT1 or DGAT2 enzymes. To determine which DGAT activity was most affected by the absence of FIT2, we treated cells with pharmacologic inhibitors specific to each DGAT enzyme and analyzed the impact on LD numbers and size 48 h after oleate addition. For wild-type cells, as expected, inhibiting both DGAT1 and DGAT2 blocked LD formation (Figure 2A). Consistent with previous findings, inhibiting either DGAT1 or DGAT2 alone did not block LD formation in wild-type cells, indicating redundancy of the enzymes in TG synthesis and LD formation (Chitraju et al., 2017). As expected, inhibiting DGAT1 alone in wild-type cells decreased LD numbers and increased the diameters of LDs, presumably through the LD expansion pathway that involves DGAT2 (Wilfling et al., 2013), and inhibiting DGAT2 resulted in increased numbers of smaller LDs, presumably via the ER-based DGAT1 pathway. The deletion of FIT2 lowered the numbers and size of LDs in all conditions, and inhibiting DGAT1 in FIT2-KO cells was sufficient to nearly completely block the formation of LDs (Figure 2A). Inhibiting DGAT2 had a much weaker effect. These results suggest FIT2 works primarily with the DGAT2 pathway.

**Figure 2:**
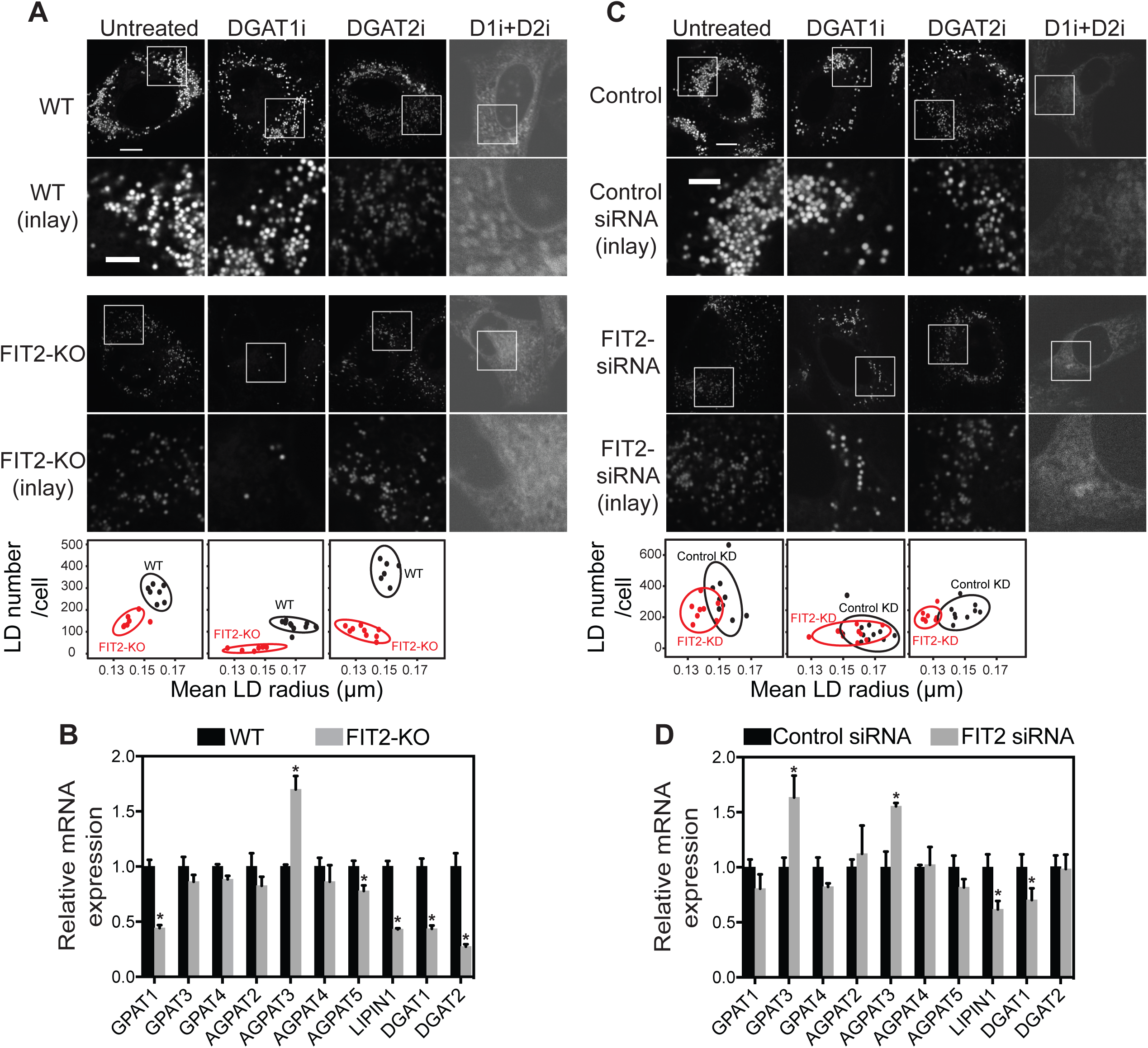
FIT2-depleted cells have a deficient DGAT2-dependent LD formation pathway. (**A**) FIT2-KO cells form predominantly DGAT1-dependent LDs. Wild-type and FIT2-KO cells were treated 2 h prior to oleic acid addition with DGAT1 and/or DGAT2 enzyme inhibitors as indicated. Untreated cells were used as control. LDs were stained with BODIPY 493/503 and cells were imaged 48 h after oleate addition. Bar, 5 µm. Average LD number and mean LD radius were measured for the indicated conditions. Each dot represents ≈17 cells. The ellipses represent a confidence interval of 90%. (**B**) Reduced mRNA expression of lipid synthesis enzymes in FIT2-KO cells. mRNA levels of genes involved in the Kennedy pathway in wild-type and FIT2-KO cells were assessed by qPCR analysis. Values represent mean ±S.D., relative to wild-type value, n=3. The figure is representative of three independent experiments (two-way ANOVA, Sidak test, \**p*<0.01). (**C**) FIT2 knock-down cells form predominantly DGAT1-dependent LDs. Wild type cells were pre-treated for 48 h with siRNA targeting FIT2 or control siRNA, and then oleic acid was added together with DGAT1 and/or DGAT2 enzyme inhibitors as indicated for another 48 h before imaging. Bar, 5 µm. Average LD number and mean LD radius were measured for the indicated conditions. Each dot represents ≈17 cells. The ellipses represent a confidence interval of 90%. (**D**) Knock-down of FIT2 results in reduced DGAT1 and LIPIN1 mRNA expression levels. mRNA levels of genes involved in the Kennedy pathway were assessed by qPCR analysis of wild-type cells pre-treated with control or FIT2 siRNA for 72 h cells. Values represent mean ± S.D. relative to wild-type value, n=3. The figure is representative of two independent experiments (two-way ANOVA, Sidak test, \**p*<0.05).

To determine how the effects of FIT2 depletion on LDs relate to changes in gene expression, we measured the mRNA expression levels of genes involved in TG synthesis by qPCR. Whereas no significant differences were found for GPAT enzymes and most of the AGPAT enzymes, the mRNA levels for LIPIN1, DGAT1, and DGAT2 in FIT2-KO cells were markedly lower (Figure 2B). Importantly, similar LD-related phenotypes were found in a second FIT2-KO cell line (referred to as clone 2), which is compound heterozygous (with one allele harboring a large deletion of FIT2 coding sequence and the second allele encoding a protein lacking a single amino acid), leading to greatly reduced expression of FIT2 (~20% of FIT2 mRNA left as determined by qPCR analysis; Figure S1A-F).

To test whether the gene expression changes found in FIT2-KO clones were due to long-term FIT2 depletion from cells, we examined gene expression after acute FIT2 knockdown (FIT2-KD) using siRNA (FIT2-siRNA). FIT2 mRNA levels were 90% lower after 72 h of siRNA treatment (Figure S2A-B). Similar to FIT2-KO cells, FIT2-KD cells formed smaller LDs, DGAT1 inhibition strongly reduced LD numbers, and DGAT2 inhibition had little effect (Figure 2C-D). As in the FIT2-KO cells, LIPIN1 and DGAT1 mRNA levels were reduced in FIT2-KD cells (Figure 2E). However, in contrast to FIT2-KO cells, DGAT2 mRNA was similar in FIT2-KD and control cells. These results support the hypothesis that FIT2 works primarily with the DGAT2 pathway, and DGAT2 downregulation found in FIT2-KO cells is due to secondary effects of longer-term FIT2 depletion.

### FIT2 deficiency impairs ER morphology

The results of FIT2 deletion on LD formation were relatively modest compared with its essential requirement in organisms, and thus, we sought to explore other ER-related phenotypes of FIT2 deficiency. Specifically, we analyzed ER structure by analyzing the distribution of ER proteins [i.e., BFP targeted to the lumen of the organelle (ss-BFP-KDEL) and the ER structural protein Sec61 tagged with GFP]. Compared with the reticular pattern of wild-type cells, FIT2-KO cells exhibited a dramatically altered ER morphology (Figure 3A). The ER was often found in clumps, suggesting aggregated membranes. Ultrastructural analyses of FIT2-KO cells by thin-section electron microscopy showed that the clumps of ER found with light microscopy were caused by abundant whorls of membranes (Figure 3B). These appeared to be ER membranes, as ribosomes were found on the membranes. ER whorls were found in 39 of 71 thin sections from FIT2 knockout cells. FIT2-KO cells also exhibited frequent occurrences of dilated ER (Figure 3C). Using lattice light-sheet microscopy of cells expressing ER-oxGFP, we found that the ER clumps in FIT2-KO cells were typically located close to the nuclear envelope and reached up to 5 μm in diameter and 20 μm^3^ in volume (Figure 3D).

**Figure 3:**
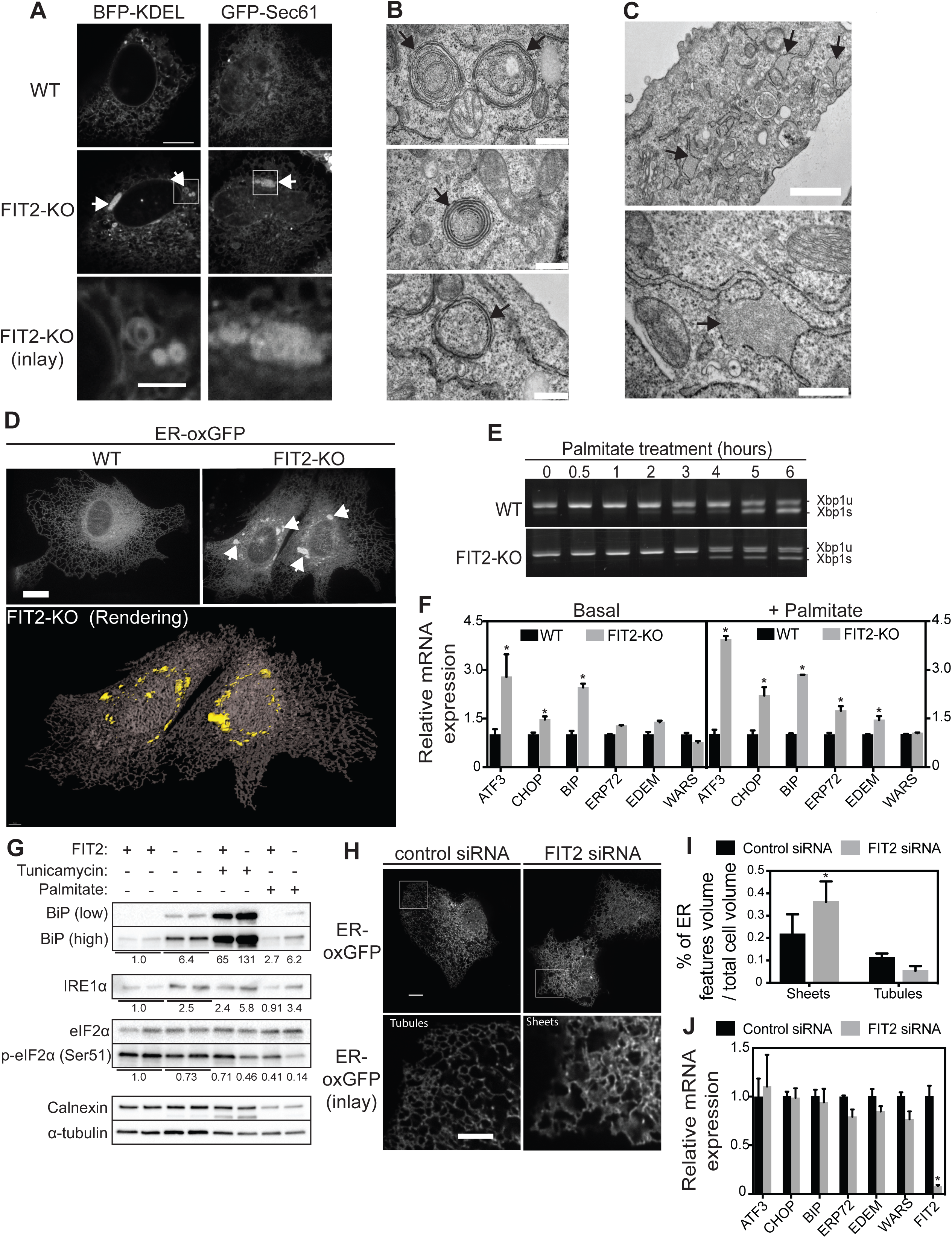
FIT2 depletion in human cells results in abnormal ER morphology and increased ER stress. (**A-D**) FIT2-deficient SUM159 cells have altered ER morphology. (A) Confocal images of wild-type and FIT2-KO SUM159 cells transiently expressing the ER marker ssBFP-KDEL or GFP-Sec61β. White arrows indicate ER aberrations. Bar, 10 µm. (B-C) Representative thin-section electron micrographs from FIT2-KO cells showing (B) ER whorls highlighted with black arrows (bar, 500 nm) and (C) ER dilation highlighted with black arrows. Bar, 1 µm (top image), 500 nm (bottom image). (D) ER whorls in FIT2-deficient cells are localized in the perinuclear environment. Representative lattice light-sheet microscopy images of wild-type and FIT2-KO cells transiently expressing oxGFP-KDEL on a plasmid. ER abnormalities are indicated with white arrows (bar, 10 µm). A rendering image performed with Imaris software shows the ER meshwork (brown) and the perinuclear localization of ER whorls (yellow shapes). (**E**) The ER stress marker Xbp1 is normally regulated in FIT2-KO cells. Xbp1 mRNA from wild-type or FIT2-KO cells, treated for the indicated times with 100 µM palmitate, was amplified by qPCR and loaded on agarose gel. Unspliced XBP1 (Xbp1u) was observed as a 152-bp band, and the spliced form (XBP1s) was observed as a 126-bp band. (**F-G**) Several ER stress markers are elevated in FIT2-KO cells. (F) qPCR analysis of the indicated genes was performed for the indicated genes on wild-type and FIT2-KO cells treated or not with 100 µM palmitate for 16 h. Values represent mean ± S.D., relative to wild-type value, n=3 (two-way ANOVA, Sidak test, \**p<*0.05). (G) Western-blot analysis of the indicated proteins was performed on wild-type and FIT2-KO cells treated or not with 100 µM palmitate for 16 h or Tunicamycin (4 µg/ml) for 14 h. Quantification of bands intensity relative to wild-type untreated is shown under each panel. (**H-I**) FIT2-KD cells have more ER sheets than control KD. (H) Confocal images of wild-type cells transiently expressing the ER marker oxGFP-KDEL treated with control siRNA or FIT2 siRNA for 72 h. An inlay is also presented highlighting the higher proportion of sheet-like structures at the cell periphery observed in FIT2 siRNA treated cells, compared to control. Bar, 10 µm. (I) Quantification of ER features (sheets and tubules) volume relative to the cell volume in cells treated with control siRNA or FIT2 siRNA for 3 days and transiently expressing the ER marker oxGFP-KDEL. Quantification is performed as described in Methods (two-way ANOVA, Bonferroni test, \**p<*0.0001). (**J**) Knock-down of FIT2 does not induce ER stress. mRNA levels of the indicated genes were assessed by qPCR analysis of wild-type cells pre-treated with control or FIT2 siRNA for 72 h. Values represent mean ± S.D., relative to wild-type value, n=3. The figure is representative of two independent experiments (two-way ANOVA, Sidak test, \**p*<0.05).

The ER whorls in FIT2-KO cells were reminiscent of membrane whorls observed during unresolved ER stress in yeast (Schuck et al., 2009). Membrane homeostasis in the ER is achieved in part via the unfolded protein response (UPR) (Ron and Walter, 2007), which is triggered by ER stress caused by the accumulation of misfolded proteins (Kozutsumi et al., 1988) or imbalanced lipids (J. Han and Kaufman, 2016; Volmer and Ron, 2015). Activation of the UPR triggers the upregulation of a large set of ER proteins, including phospholipid synthesis enzymes that provide more membrane lipids (Sriburi, 2004; Travers et al., 2000). In FIT2-KO cells, we found no evidence of increased activation of Xbp1 splicing (a marker of ER stress), either at baseline or after palmitate addition (to exacerbate lipid-induced ER stress) (Figure 3E). However, we found a modest upregulation of mRNA levels of other ER stress markers, such as ATF3, BIP, and CHOP, in FIT2-KO cells cultured under basal conditions that were further increased with palmitate addition (Figure 3F). The activation of the ER stress pathway in FIT2-KO cells was confirmed by immunoblotting, which showed increased levels of BIP and Ire1α. In contrast, we did not detect activation of eIF2α, suggesting that the PERK pathway is not constitutively activated in FIT2-KO cells (Figure 3G). ER morphology was also affected in acutely FIT2-depleted (FIT2-KD) cells, which exhibited a higher proportion of apparent ER sheets in the cell periphery (Figure 3H-I; Figure S2C). However, activation of ER stress was not detected in FIT2-KD cells (Figure 3J), possibly due the lack of activation at the time point sampled.

In contrast to the prominent effects of FIT2 deficiency on the ER, no differences were found in the appearance of the Golgi apparatus, lysosomes, or peroxisomes (Figure S3).

### FIT2 is a lipid phosphate phosphatase

Our findings indicated that FIT2 is crucial for normal ER homeostasis. The presence of orthologs in most eukaryotes (Kadereit et al., 2008), and the severe phenotypes of FIT2 deletion (Choudhary et al., 2015; Goh et al., 2015), suggested that FIT2 carries out an important evolutionarily conserved function. Analysis of FIT2 amino acid sequences among species revealed that the overall architecture of FIT2 and evolutionarily homologous proteins (e.g., yeast Scs3) is similar to lipid phosphate phosphatases (LPP enzymes; Figure 4A). LPP enzymes include a large number of proteins that hydrolyze lipid phosphates, such as phosphatidic acid and lysophosphatidic acid (Dillon et al., 1997; Toke et al., 1998a; 1998b; Waggoner et al., 1996; 1995), but also include transferases such as sphingomyelin synthase (SMS; Huitema et al., 2004). Importantly, the catalytic residues in LPP enzymes are conserved in FIT2 and its yeast homologue Scs3 (Figure 4B). In particular, this includes two histidines contained in two catalytic motifs, designated C2 and C3. The C3 histidine (H214 for FIT2 and H350 for Scs3) acts on the phosphorus of the phospholipid as a nucleophile for the formation of a phospho-histidine intermediate. The C2 histidine (H155 for FIT2 and H235 for Scs3) is involved in phosphate-bond breaking to release the de-phosphorylated substrate. There is also a C1 domain (KXXXXXXRP) in most LPP enzymes that is thought to play a role in substrate recognition (Sigal et al., 2005), but this domain does not appear to be present within the FIT2 family (Figure 4C). Mapping these potentially catalytic residues on the current model of FIT2’s membrane topology (Gross et al., 2010) revealed that the conserved catalytic residues are predicted to be positioned at the interface between the inner ER membrane leaflet and the ER lumen (Figure 4D).

**Figure 4:**
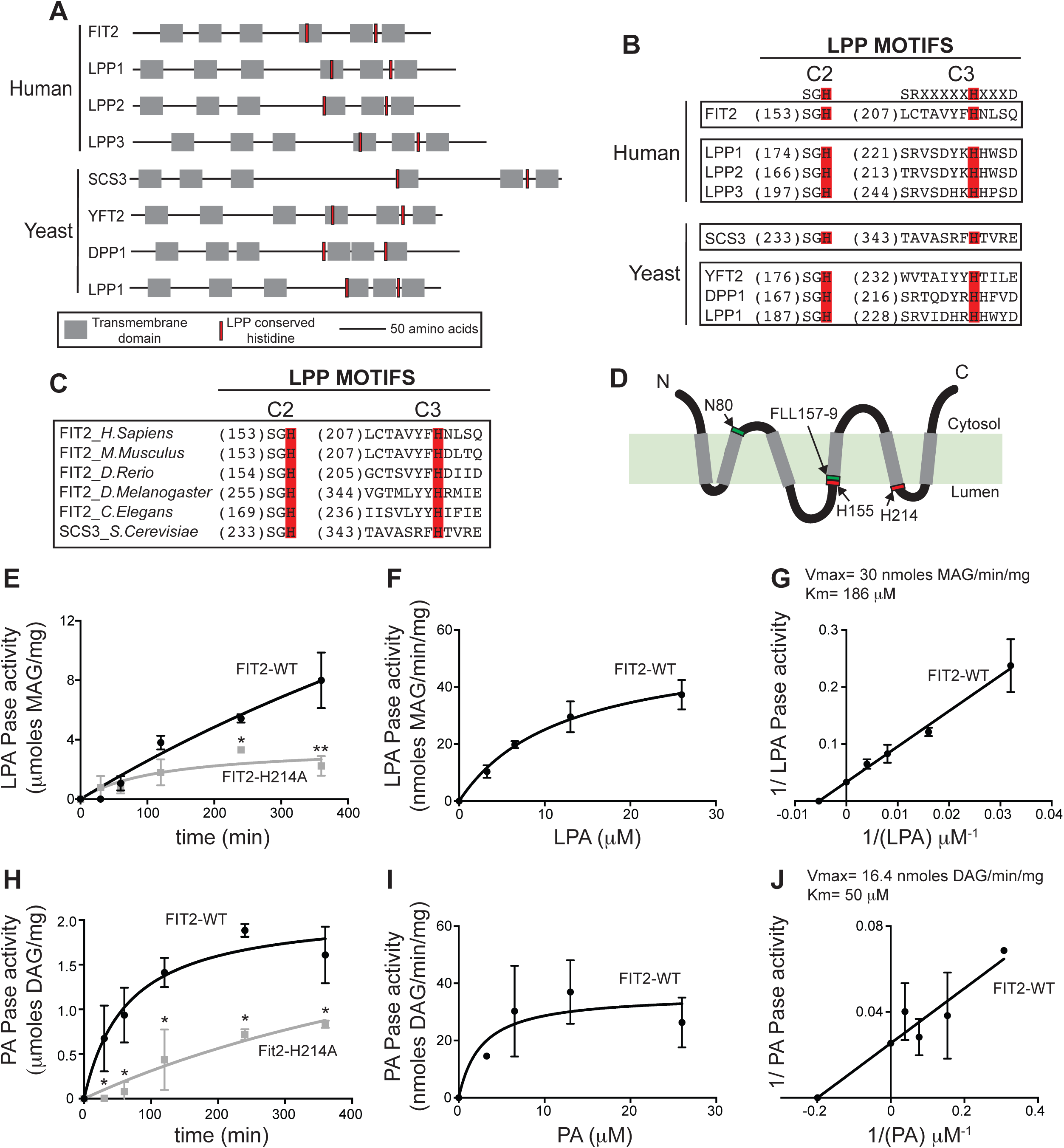
Recombinant human FIT2 possesses lipid phosphate phosphatase activity. (**A**) FIT2 overall topology (six transmembrane domains) and the positions of C2 and C3 histidines are similar to those of other LPP enzymes (Lpp1, Lpp2 and Lpp3). Also shown are Scs3, its homolog Yft2 and some yeast LPP members (Dpp1 and Lpp1). (**B**) FIT2 shares conserved motifs with LPP enzymes. FIT2 and its yeast homolog Scs3 have two conserved LPP motifs. C2 and C3 motifs of FIT2 and Scs3 were aligned using human and yeast LPP members. The conserved catalytic histidines are highlighted in red. (**C**) C2 and C3 catalytic histidines (highlighted in red) are evolutionary conserved within FIT2 sequences. (**D**) Based on topology analysis (Gross et al., 2010), FIT2 N- and C-termini are predicted to be cytosolic, and the C2 and C3 motifs are predicted to be oriented toward the ER lumen. Transmembrane domains are represented as gray bars. Red boxes represent the C2 and C3 catalytic histidines. The position of the amino acids that affect diacylglycerol and triacylglycerol binding (N80 and FLL157-159) (Gross et al., 2011) are also shown in a green boxes. (**E-F-G**) FIT2 has lysophosphatidic acid (LPA) phosphatase activity. LPA phosphatase activity was measured as the formation of monoacylglycerol (MAG) on TLC as described in Methods. (E) Time-dependent LPA phosphatase assay: 20 nmoles of LPA was incubated with 100 ng of wild-type FIT2, FIT2-H214A. The amount of MAG produced at each time point was quantified by scintillation counting from triplicate enzyme determinations, and mean values ±SD were plotted over time. (two-way ANOVA, Sidak test, \**p*<0.05; \*\**p*<0.0001). (F) Substrate-dependent LPA phosphatase assay. The indicated concentration of LPA was incubated for 1 h with 100 ng of wild-type FIT2. The amount of MAG produced for each LPA concentration was quantified by scintillation counting from triplicate enzyme determinations, and mean values ±SD were plotted over LPA concentration. (G) Same data as in (**F**) were used to make Lineweaver-Burk plot and to determine Vmax and Km. (**H-I-J**) FIT2 has phosphatidic acid (PA) phosphatase activity. (H) 20 nmoles of PA was incubated with 100 ng of wild-type FIT2, FIT2-H214A. The amount of DAG produced at each time point was quantified by scintillation counting from triplicate enzyme determinations, and mean values ±SD were plotted over time. (two-way ANOVA, Sidak test, \**p*<0.05). (I) Substrate-dependent PA phosphatase assay. The indicated concentration of PA was incubated for 1 h with 100 ng of wild-type FIT2. The amount of DAG produced for each PA concentration was quantified by scintillation counting from triplicate enzyme determinations, and mean values ±SD were plotted over PA concentration. (J) Same data as in (I) were used to make Lineweaver-Burk plot and to determine Vmax and Km.

To directly test whether FIT2 has LPP activity, we purified human wild-type protein, as well as a mutant form of the protein in which the key catalytic C3 histidine (H214) was replaced with an alanine residue (FIT2 H214A; Figure S4A-C). When incubated *in vitro* with either phosphatidic acid or lysophosphatidic acid substrates, the wild-type protein increased the rate of lipid phosphate hydrolysis. This activity was much lower when substrates were incubated with similar amounts of FIT2 H214A (Figure 4E and 4H). Under our assay conditions, the specific activity of wild-type FIT2 towards lysophosphatidic acid showed an apparent Km = 186 µM and a Vmax = 30 nmoles MAG/min/mg. (Figure 4F-G). The specific activity of FIT2 towards phosphatidic acid showed an apparent Km = 50 µM and a Vmax = 16.4 nmoles DAG/min/mg (Figure 4I-J). These data show that FIT2 has lipid phosphatase activity on several known LPP substrates *in vitro*.

To determine if FIT2 has activity for a phospholipid substrate with a head group, we tested whether FIT2 could catalyze the hydrolysis of lysophosphatidylcholine (LPC) to form MAG. In contrast to control activity catalyzed by phospholipase C, FIT2 did not exhibit LPC hydrolase activity (Figure S4D).

### FIT2 catalytic residues are required for ER maintenance in *Saccharomyces cerevisiae*

We next tested whether the LPP activity of FIT2 is crucial for its function *in vivo*. In agreement with our findings in human FIT2-KO cells, yeast *SCS3* was required for maintaining a normal ER, and in the absence of Scs3, abundant clumps of ER membrane were found (detected by the ER marker signal sequence-RFP-HDEL) (Figure 5A). This phenotype was complemented by expressing wild-type Scs3, but not by expressing mutant forms of the protein [with alanine mutations of the catalytic residues (Scs3 H235A, Scs3 H350A)] (Figure 5B). The ER phenotype was also complemented by expressing human FIT2 in *scs3*Δ cells, but not by expressing catalytically defective forms of human FIT2 (H155A and FIT2 H214A) (Figure 5C).

**Figure 5:**
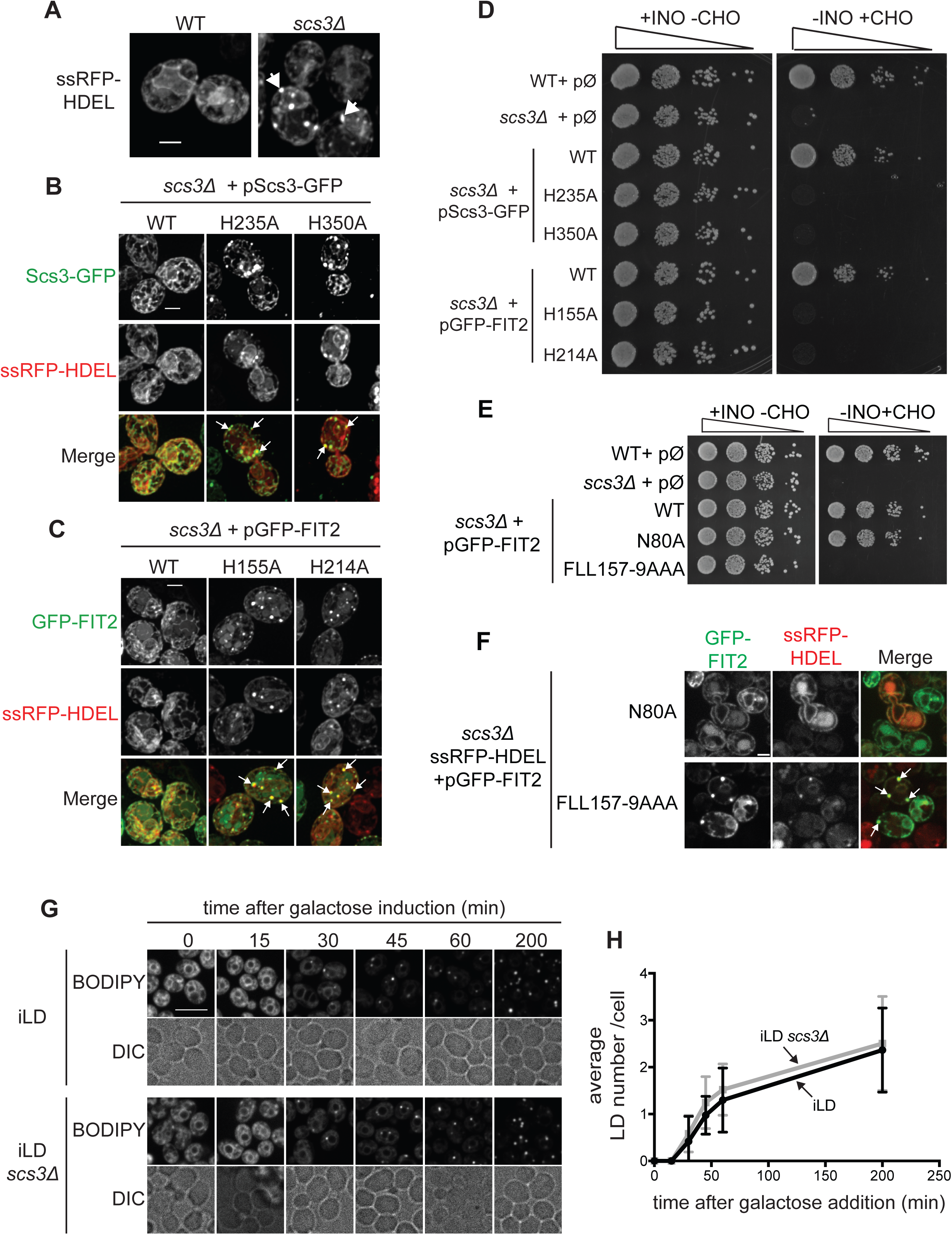
Mutations of putative catalytic LPP residues compromise Scs3/FIT2 *in vivo* function in yeast. (**A**) *SCS3* deletion triggers the formation of ER patches. Max-Z projection of representative deconvolved confocal images from wild-type and *scs3*∆ cells expressing the ER marker ssRFP-HDEL on a plasmid; White arrows highlights ER patches. Scale: 2 µm. (**B**) Reintroduction of Scs3-GFP mutated at C2 or C3 histidine does not restore normal ER shape in *scs3*∆. Subcellular localization of Scs3-GFP wild-type or mutated on C2 or C3 histidine to alanine (H235A and H350A, respectively) was assessed in *scs3*∆ cells. Max-Z projection of representative deconvolved confocal images is presented. Impact on ER shape was addressed with ssRFP-HDEL marker; white arrows highlight ER patches. Scale: 2 µm. (**C**) Reintroduction of GFP-FIT2 mutated at C2 or C3 histidine does not restore normal ER shape in *scs3*∆. Subcellular localization of GFP-FIT2 wild-type or mutated on C2 or C3 histidine to alanine (H155A and H214A, respectively) was assessed in *scs3*∆ cells. Max-Z projection of representative deconvolved confocal images is presented. Impact on ER shape was addressed with ssRFP-HDEL marker; White arrows highlights ER patches. Scale: 2 µm. (**D**) C2 or C3 histidine mutations of Scs3 or FIT2 do not rescue *scs3*∆ inositol auxotrophy. wild-type cells, *scs3*∆ cells, and *scs3*∆ cells expressing GFP-tagged Scs3 or FIT2, wild-type or mutated on C2 histidine to alanine (H235A for Scs3, H155A for FIT2) or C3 histidine to alanine (H350A for Scs3, H214A for FIT2) on a plasmid. Cells were serially diluted and spotted on complete solid medium (+INO-CHO) or inositol-depleted (-INO+CHO). To strengthen the growth defect as described in (Villa-García et al., 2010), inositol-deprived medium was supplemented with choline (+CHO), and cells were grown at 37° C. The plates were imaged after 4–5 days. (**E-F**) Mutant form of FIT2 having high DAG/TG binding affinity is not functional. (E) N-ter GFP-tagged mutant versions of FIT2 increasing DAG/TG affinity (FLL157-9>AAA) or decreasing affinity (N80A) (Gross et al., 2011) were expressed on a plasmid in *scs3*∆ cells and plated on complete (+INO-CHO) or inositol-deprived medium (-INO+CHO). (F) Subcellular localization of the constructs in *scs3*∆ cells used in (E) was assessed by confocal imaging. Impact on ER shape was addressed with ssRFP-HDEL ER marker; white arrows highlights ER patches. Bar, 2 µm. (**G**) Deletion of *SCS3* does not affect LD formation in yeast. Representative confocal images of inducible LD strain deleted for *SCS3* (iLD *scs3*∆) or not (iLD) after overnight growth in raffinose (repressed LD condition) and after galactose addition (induced LD condition) at the indicated times. LDs were stained with BODIPY 493/503. Scale: 5 µm. (**H**) Average LD number per cell ±SD was measured over time, based on images presented in (E) (n=100).

Consistent with a role in lipid metabolism, Scs3 is required for inositol auxotrophy (Hosaka et al., 1994). Consequently, *scs3*Δ cells fail to grow on plates with medium lacking inositol (Figure 5D). This phenotype was complemented by introducing wild-type Scs3 or human FIT2, but not with mutant forms of these enzymes (Scs3 H235A or Scs3 H350A; FIT2 H155A or FIT2 H214A) (Figure 5D). Also, introduction of either Scs3 histidine mutant into a wild-type strain did not induce the formation of ER clumps nor affect inositol auxotrophy (Figure S5B-C), indicating that expression of those mutants does not have dominant-negative effects. These data indicate that the LPP catalytic residues and likely the FIT2 lipid phosphate phosphatase activity are required for its ER-related function in yeast.

In addition, we investigated mutants previously found to be deficient in binding neutral lipids TG (Gross et al., 2011). These include FLL157-159>AAA, which displays higher affinity for DAG/TG and considered as a gain-of-function mutant, and N80>A which had less affinity for neutral lipids and was characterized as a loss-of-function mutant (Gross et al., 2011). Surprisingly, we found that expression of either WT or the FIT2-N80>A mutant in *scs3*Δ cells rescued both the inositol auxotrophy and the ER morphology phenotypes, whereas expression of FLL157-159>AAA did not (Figure 5E-F). These data show that FIT2-FLL157-159>AAA is not functional. Since these mutations are close to the putative catalytic residue H155, one possible explanation to these findings could be that higher affinity for DAG found with the FLL157-159>AAA mutation blocks the catalytic site of FIT2, acting as product-inhibition of the enzyme.

We also tested *scs3*Δ cells for LD phenotypes. In yeast, four genes encode the enzymes involved in neutral lipid synthesis (Dahlqvist et al., 2000; Oelkers et al., 2002; 2000; Sorger and Daum, 2002): *DGA1* and *LRO1* for TG synthesis and *ARE1* and *ARE2* primarily for sterol ester synthesis. We used a yeast strain in which *LRO1* and *ARE1* were deleted and where *DGA1* and *ARE2* were placed under a galactose-inducible promoter. This strain, designated as “induced LD” (iLD) enabled us to induce LD formation by switching cells from raffinose-to galactose-containing medium (similar to strain used in (Cartwright et al., 2014). After induction, we found the formation of LDs, assessed by BODIPY staining, at ~30 min (Figure 5G), and the additional deletion of *SCS3* did not affect LD formation (Figure 5G-H).

### FIT2 catalytic residues are required for normal ER in mammalian cells

To further test the hypothesis that FIT2 acts as a lipid phosphate phosphatase affecting ER homeostasis in mammalian cells, we expressed wild-type or mutants of the catalytic residues in the FIT2-KO SUM159 cells. Expression of wild-type FIT2, but not catalytic mutant forms, completely reversed the ER morphology phenotype associated with FIT2 depletion (Figure 6A-B).

**Figure 6:**
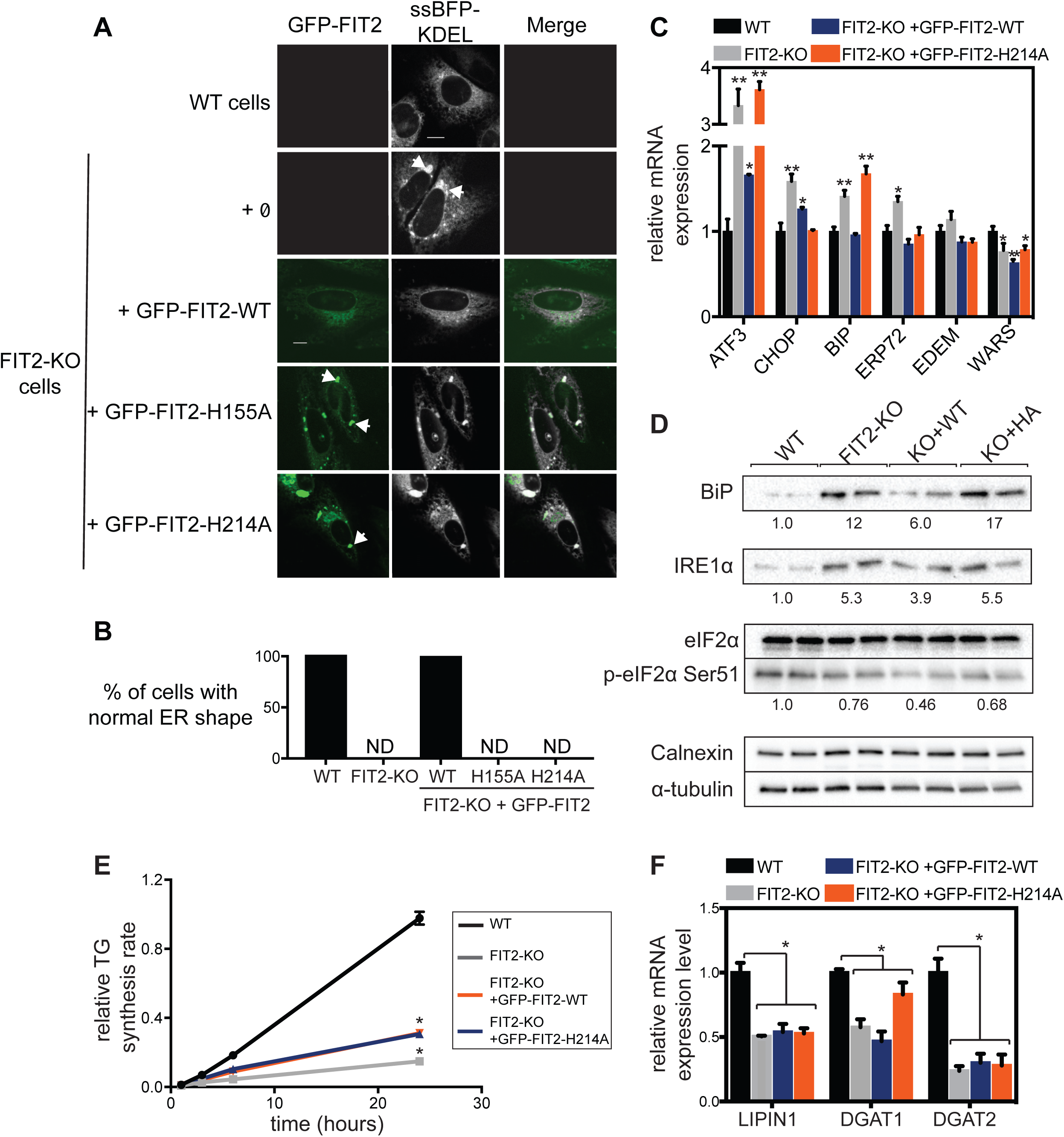
Putative catalytic residues in FIT2 are required for normal ER morphology in human cells. (**A-B**) Reintroduction of GFP-FIT2 mutated at LPP conserved histidines does not restore normal ER shape in FIT2-KO cells. (A) Representative confocal images of wild-type cells, FIT2-KO cells transiently co-expressing ER marker ssBFP-KDEL and GFP-FIT2 wild-type or C2 (H155A) or C3 histidine mutants (H214A). White arrows highlight ER patches. Scale: 10 µm. (B) Quantification of the percentage of rescued ER shape in FIT2-KO cells expressing wild-type GFP-FIT2, GFP-FIT2-H155A or GFP-FIT2-H214A shown in (A) (n=50). ND: Not detected. (**C-D**) Reintroduction of GFP-FIT2 mutated at H214A does not restore normal ER stress markers expression. (C) qPCR analysis of the indicated genes (values represent mean ±S.D., n=3, representative of three independent experiments; two-way ANOVA, Sidak test, \**p*<0.01; \*\**p*<0.001) and (D) Western-blot analysis of ER stress proteins in wild-type cells, FIT2-KO cells and FIT2-KO cells stably expressing either wild-type GFP-FIT2 (KO+WT) or GFP-FIT2-H214A (KO+HA). Quantification of bands intensity relative to wild-type untreated is shown under each panel. (**E**) Reintroduction of wild-type GFP-FIT2 or H155A does not restore normal TG synthesis rate in FIT2-KO cells. Cells were pulse-labeled with [^14^C]-oleic acid, and incorporation into TG was measured over time during oleate treatment by TLC. Mean value of TG band intensity was plotted over time and quantified from three independent measurements. Values were calculated relative to 24 h time point value of wild-type cells (two-way ANOVA, Sidak test, \**p*<0.01). (**F**) Reintroduction of wild-type GFP-FIT2 or H214A does not restore normal expression of neutral lipid synthesis genes in FIT2-KO cells. qPCR analysis of the indicated genes was performed on wild-type cells, FIT2-KO cells and FIT2-KO cells stably expressing either wild-type GFP-FIT2 or GFP-FIT2-H214A. Values represent mean ±S.D., n=3. The figure is representative of three independent experiments (two-way ANOVA, Sidak test, \**p*<0.01).

To test if the rescue in morphology correlated with normalization of ER stress markers, we stably reintroduced wild-type or catalytic mutant FIT2 into FIT2-KO cells with the AAVS1 “safe harbor” targeting system (System Biosciences; Figure S6). Adding back wild-type FIT2 restored normal mRNA levels of all the elevated ER stress markers of FIT2-KO cells (Figure 6C). In contrast, adding back catalytic-mutant FIT2 did not restore normal levels of Atf3 and BIP. This was further confirmed by immunoblot analysis of BIP and Ire1α. eIF2α showed no activation in any strains (Figure 6D). Surprisingly, changes in gene expression of lipid synthesis genes found in FIT2-KO cells (Figure 2C) were not restored by stable reintegration of wild-type FIT2 or FIT2-H214A (Figure 6F), nor were TG synthesis rates normalized (Figure 6E). These findings further support the hypothesis that these changes in gene expression occur as a secondary phenotype of long-term FIT2 depletion in cells.

## Discussion

We show here that FIT2, a protein previously identified as important for LD formation (Choudhary et al., 2015; Kadereit et al., 2008), is an enzyme, with purified recombinant protein exhibiting LPP activity towards two known LPP substrates, LPA and PA. Based on sequence homology with other LPP family members, we find that the predicted catalytic histidine residues of FIT2 are crucial for its enzymatic activity, and these residues are required for its biological function in cells, with mutant forms of the enzyme being unable to rescue cellular phenotypes. Taken together, our principal findings indicate that FIT2 is an ER-localized LPP that is of crucial importance for maintaining a normal ER, likely by maintaining phospholipid balance. Another recent report (Hayes et al., 2018) suggesting that FIT2 is an LPP enzyme, based on sequence analyses, is consistent with this conclusion.

It is currently unclear whether PA and LPA are *in vivo* substrates for FIT2, or alternatively if the enzyme possesses other LPP activities. Consistent with the possibility that PA is an important *in vivo* substrate, cells lacking FIT2 that were cultured with oleate accumulated PA and had lower flux of oleate tracer into other phospholipids and neutral lipids. However, these changes were only present under conditions of oleate loading, and at steady state, we found no differences in cellular lipids with FIT2 depletion, possibly because there are compensatory changes that restore lipid homeostasis unless the cells are stressed with excess lipids. Moreover, it remains possible that Fit2 dephosphorylates other substrates or acts as a transferase. For instance, homology searches using the HHPRED algorithm (Alva et al., 2016) identify the strongest similarities of human Fit2 with the bacterial lipid phosphatase PgpB, which cleaves phosphatidylglycerol phosphate.

Our studies, together with the topology of FIT2 (Gross et al., 2010), predict that the active site of FIT2 may reside on the luminal surface of the ER. Thus, FIT2 is predicted to act on the luminal side of the ER, unlike PAH1/lipin enzymes which provide much of the ER PA hydrolase activity on the cytosolic face of the ER (G. S. Han et al., 2006). Based on the available data, we hypothesize that FIT2 activity provides a neutral lipid product (e.g., DAG or MAG) that flips rapidly across the bilayer to provide a means to balance phospholipids in the two leaflets of the ER membrane. This may be particularly important when LDs are formed and bud toward the cytosolic face, increasing the requirement for phospholipids, such as PC, to move from the cytosolic face of the ER bilayer to the LD monolayer to coat the expanding droplet. The demand for PC during LD formation is considerable, considering PC levels can increase more than two-fold in cells that are cultured in oleate to stimulate LD formation (Krahmer et al., 2011). Mechanisms that facilitate balance of phospholipids between the ER membrane leaflets may be particularly important during such situations.

A major consequence of FIT2 deficiency in our studies was abnormalities in ER morphology, along with activation of ER stress pathways to restore homeostasis. Consistent with a role of FIT2 in maintaining ER structure, the yeast atlastin ortholog *SEY1*, encoding a key factor in ER shaping, is the strongest negative genetic interactor with *scs3Δ* (Surma et al., 2013).

In mammalian cells, FIT2 knock-down cells displayed increased amounts of ER sheets in the cell periphery, and FIT2-KO cells showed even more severe ER aberrations such as clumps and whorls, suggesting a correlation between the severity of the ER defect and FIT2 expression level. ER membrane whorls in cells have been found in different situations, such as unresolved ER stress (Schuck et al., 2009), PC deficiency (Testerink et al., 2009), fatty acids saturation (Surma et al., 2013), drug treatment (Feldman et al., 1981), or overexpression of ER resident proteins (Chin et al., 1982; Gong et al., 1996; Koning et al., 1996; Snapp et al., 2003; Wright et al., 1988), and thus the occurrence of whorls in FIT2-deficient cells is consistent with abnormalities in the ER phospholipids. Moreover, the ER morphology changes with FIT2 deletion are consistent with a role for FIT2 in maintaining the balance between phospholipids in the two leaflets of the ER. Whereas membrane sheets have the same amount of phospholipids in both leaflets, highly curved membranes in ER tubules require more lipids on the cytoplasmic leaflet than the luminal leaflet, and thus FIT2 deficiency would be predicted in this model to favor sheets over tubules.

Our findings suggest that alterations in LD formation from the ER with FIT2 deletion are likely secondary effects to changes in ER lipids. In mammalian cells, we found reduced numbers of LDs were formed for a given amount of TG synthesis, suggesting that LD formation is less efficient with FIT2 deletion. This conclusion would be predicted from the hypothesis that FIT2 maintains proper balance of phospholipids in the ER leaflets. Also, we found that a correlation of FIT2 activity predominantly with DGAT2 pathway of expanding LD of formation (Wilfling et al., 2013). This pathway is prominent when the need for TG storage and LD formation is maximal, consistent with FIT2 being required during lipid overload. Interestingly, we found no detectable effects on LD formation in *scs3*Δ cells in yeast perhaps because yeast generate so few LDs and phospholipids on the cytosolic leaflet are not as limiting.

The accumulating evidence suggests a model in which FIT2 LPP activity on the luminal face of the ER, is required to maintain normal ER phospholipid distribution. In basal conditions, the loss of FIT2 is largely compensated by activation of ER stress pathways, and restoration of cellular lipid levels, although cells exhibit signs of FIT2 deficiency, such as membrane whorls and more ER sheets. During times of increased needs for phospholipids, e.g., during oleate loading and LD formation, deficiency of FIT2 impacts LD formation, which is highly sensitive to changes in membrane composition (Ben M’barek et al., 2017; Choudhary et al., 2018) and, we hypothesize, requirements for more phospholipids on the cytosolic face. Therefore, in the absence of FIT2, LD formation is less efficient. We speculate that demands for FIT2 activity to maintain ER lipid balance in specific tissues results in the severe consequences found in FIT2 deficiency, such as lethality in model organism knockouts (Choudhary et al., 2015; Goh et al., 2015) and syndromes of deafness and myotonia in humans (Seco et al., 2016).

## Acknowledgments

We thank members of the lab for technical assistance and helpful comments on the manuscript, David Silver, Sebastien Léon, and George Carman for reagents, Tim Levine for helpful discussions, Gary Howard for editorial assistance. MB is a recipient of a fellowship from the Jane Coffin Childs Memorial Fund for Medical Research. NM is a recipient of a fellowship from Marie Sklodowska-Curie. This work was supported by NIH grant 5R01GM124348 (to R.F.) and R01GM097194 (to T.C.W.). TCW is a HHMI investigator.

## Supplementary figures

**Figure S1. Related to Figure 1-2:**
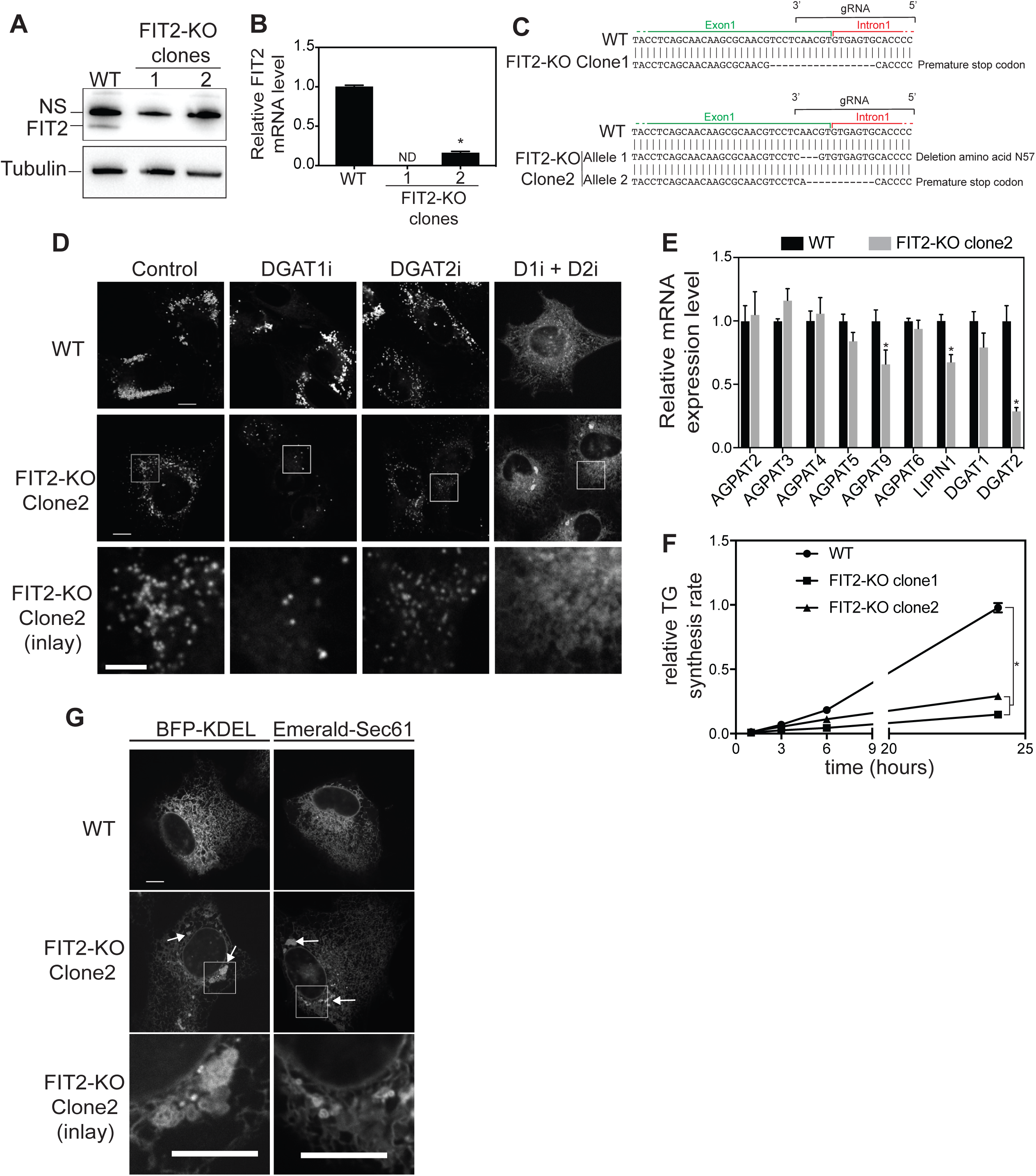
Characterization of FIT2-KO SUM159 clones. (**A**) No detectable FIT2 protein was observed for both FIT2-KO clones. Western blot on wild-type, FIT2-KO clone1 (used in the entire study), and FIT2-KO clone2 crude extracts revealed with anti-FIT2 antibody. (**B**) 20% expression level of FIT2 was detectable in FIT2-KO clone2 cells, 0% for clone1. FIT2 mRNA level analysis by qPCR in wild-type and FIT2-KO clones. Values represent mean ±S.D. relative to wild-type cells level, n=3 (two-way ANOVA, Sidak test, \**p*<0.001). ND: Not detected. (**C**) Sequence of genome edited region in FIT2-KO clone1 and clone2 cells. FIT2-KO clone1 contains a 17-bp deletion located at the end of exon 1 in FIT2 locus, leading to a frame shift. FIT2-KO clone2 contains an 11-bp deletion located at the end of exon 1 in FIT2 locus, leading to a frame shift. In the other allele, there is a 3-bp deletion, leading to the deletion of an asparagine in position 57. gRNA position in wild-type genome is indicated. (**D**) FIT2-KO clone2 cells form only DGAT1-dependent LDs. Wild-type or FIT2-KO clone2 cells were pre-treated with DGAT1 and/or DGAT2 enzyme inhibitors as indicated for 2 h, and then oleic acid was added for another 48 h before imaging. Representative confocal images are presented. Bar, 10 µm. (**E**) FIT2-KO clone2 displays an alteration of lipid synthesis enzymes expression. mRNA levels of genes involved in the Kennedy pathway were assessed by qPCR analysis of wild-type and FIT2-KO clone2 cells. Values represent mean ±S.D. relative to wild-type cells level, n=3 (two-way ANOVA, Sidak test, \**p*<0.05). (**F**) FIT2-KO clone2 displays a TG synthesis defect. Wild-type and FIT2-KO clone2 cells were pulse-labeled with [^14^C]-oleic acid, and incorporation into TG was measured over time during oleate treatment by TLC. Mean value of TG band intensity was plotted over time and quantified from three independent measurements. Values were calculated relative to 24 h time point value of wild-type cells (two-way ANOVA, Sidak test, \**p*<0.01). TG synthesis rate of FIT2-KO clone1 is also reported as a comparison. (**G**) ER morphology is affected in FIT2-KO clone2. Confocal images of wild-type and FIT2-KO clone2 cells transiently expressing the ER marker ssBFP-KDEL or GFP-Sec61β. White arrows indicate ER aberrations. Bar, 10 µm.

**Figure S2. Related to Figure 2 and 3:**
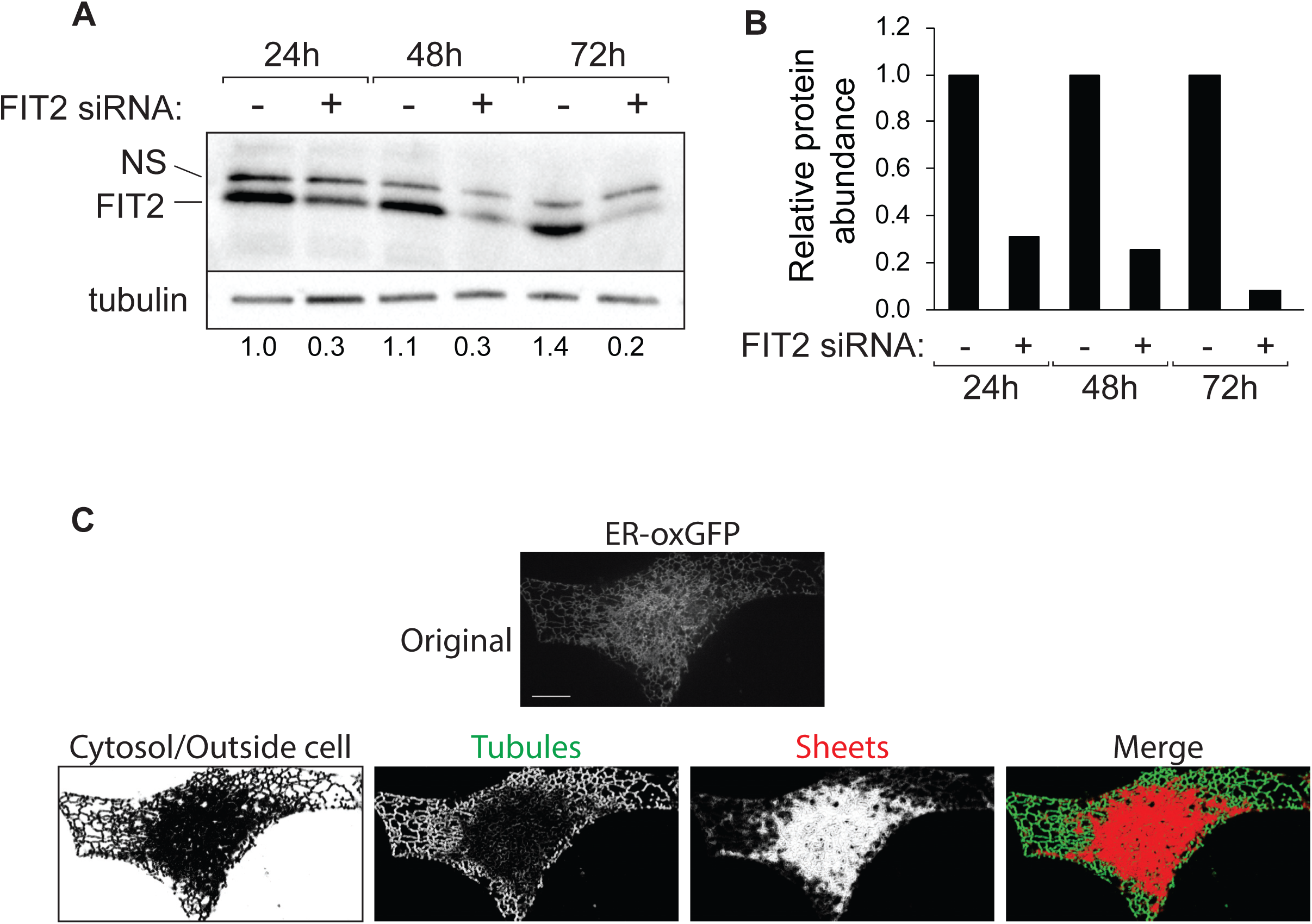
FIT2 knock-down efficiency and ER features quantification. (**A**). Cells treated with FIT2-siRNA treatment for 72 h results in 80% FIT2 protein reduction. Cells treated with control or FIT2 siRNA for the indicated time and FIT2 knock-down efficiency was followed by western blotting with anti-FIT2 antibody. Tubulin is used as loading control. Quantification of bands intensity relative to control siRNA at 24 h is shown under tubulin panel (**B**) Quantification of FIT2 protein abundance based on western blot shown in (A) was plotted relative to control siRNA for each time points. (**C**) Segmentation of ER feature using machine learning. Cells treated with control siRNA for 72 h were transiently expressing the ER marker ER-oxGFP on a plasmid. One confocal image of a Z-stack is presented (original image). As detailed in the methods, machine learning was used to segment the ER features (sheets, tubules and outside of the cell/cytosol). Bar, 10 µm.

**Figure S3. Related to Figure 3:**
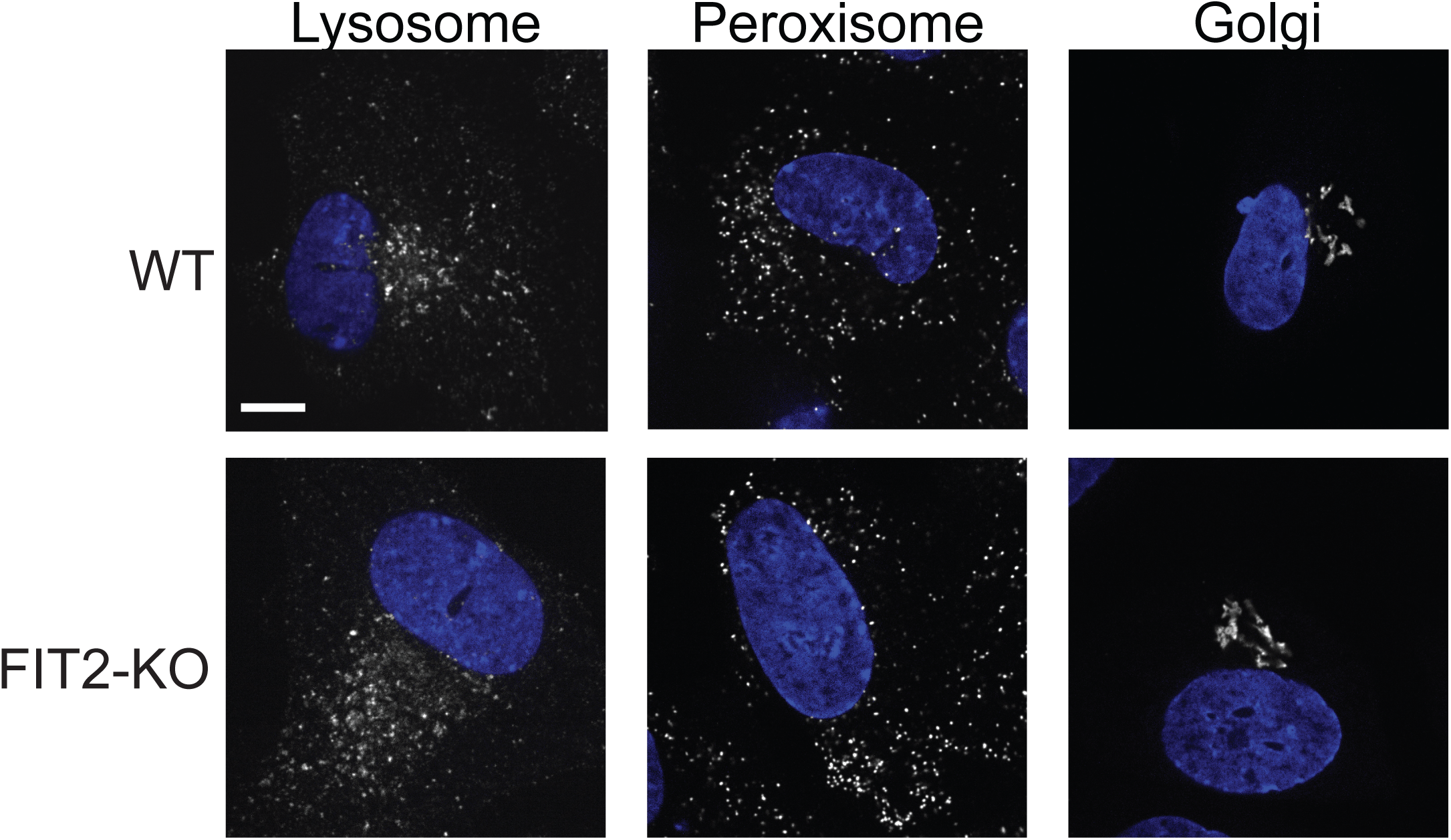
FIT2 depletion does not affect the morphology of other organelles. Representative confocal images of immunofluorescence using antibodies targeting markers of Golgi apparatus (anti-GM130), Peroxisome (anti-catalase), Lysosome (anti-LAMP1) in wild-type and FIT2-KO SUM159 cells. Bar, 10 µm.

**Figure S4. Related to Figure 4:**
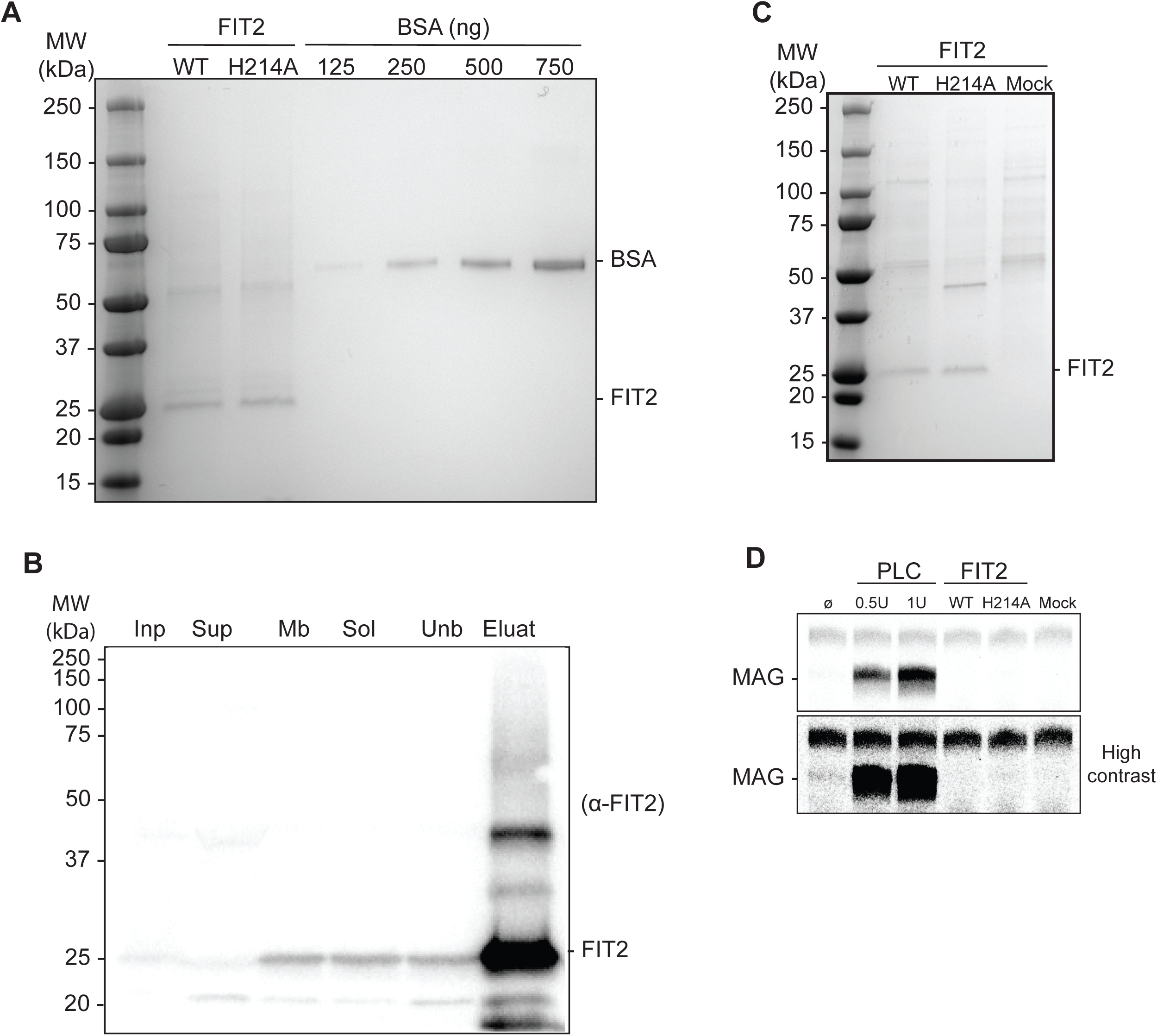
Purification of FIT2 recombinant protein. (**A**) FIT2 was expressed and purified from *S. cerevisiae* as described in experimental procedures. Purification of recombinant StrepII-tagged FIT2 protein wild-type or C3 Histidine mutant (H214A) was assessed on SDS-polyacrylamide gel stained with Coomassie Brilliant blue. Protein concentration was calculated by comparing band intensity with bovine serum albumin (BSA) standards. (**B**) Western blot of the purification steps was performed using anti-FIT2 antibody. FIT2 protein is observed at the expected size (22 KDa). *Inp,* Input; *Sup*, Supernatant (cytosolic protein); *Mb*, Total Membrane fraction; *Solub*, Solubilized fraction; *Unb*, Unbound fraction after incubation with Streptactin beads. (**C**) An additional purification was also performed with *scs3*∆ cells extract not expressing FIT2 (Mock) to assess the contribution of contaminant proteins in the enzymatic assays. (**D**) FIT2 does not have lysophosphatidylcholine (LPC) lipase activity. LPC lipase activity was determined by following the formation of MAG on TLC. 20 nmoles of LPC was incubated with 100 ng FIT2 wild-type (WT), FIT2-H214A, Mock extract or no protein (ø) for 1 h. 0.5 or 1 unit of phospholipase C (PLC) were used as positive controls.

**Figure S5. Related to Figure 5:**
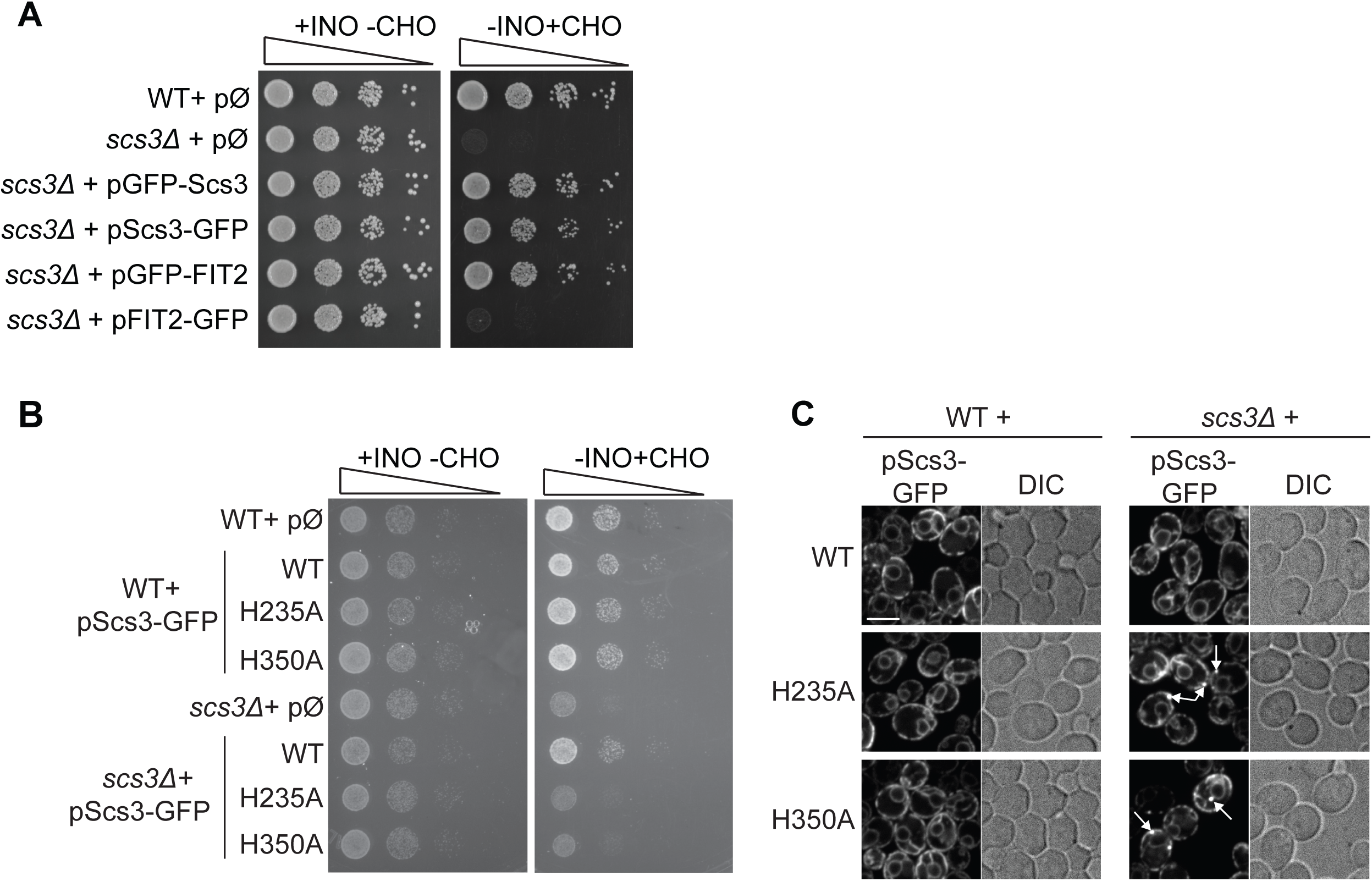
Yeast functional assays. (**A**) Both N- and C-ter GFP-tagged Scs3 are functional, whereas only N-ter GFP-tagged FIT2 is functional. N- and C-ter GFP-tagged versions of Scs3 and FIT2 expressed on a plasmid in *scs3*∆ cells were grown on complete (+INO-CHO) or inositol-deprived medium (-INO+CHO). (**B-C**) Scs3 LPP mutants do not have dominant-negative effects. (B) Wild-type or LPP mutant forms of Scs3 expressed on plasmid in wild-type cells were grown on complete (+INO-CHO) or inositol-deprived medium (-INO+CHO) for 2 days, and (C) their subcellular localizations were assessed by confocal fluorescence microscopy. Localization of the same constructs in *scs3*∆ cells is also presented. Presence of ER whorls is indicated with white arrows. Bar, 5 µm.

**Figure S6. Related to Figure 6:**
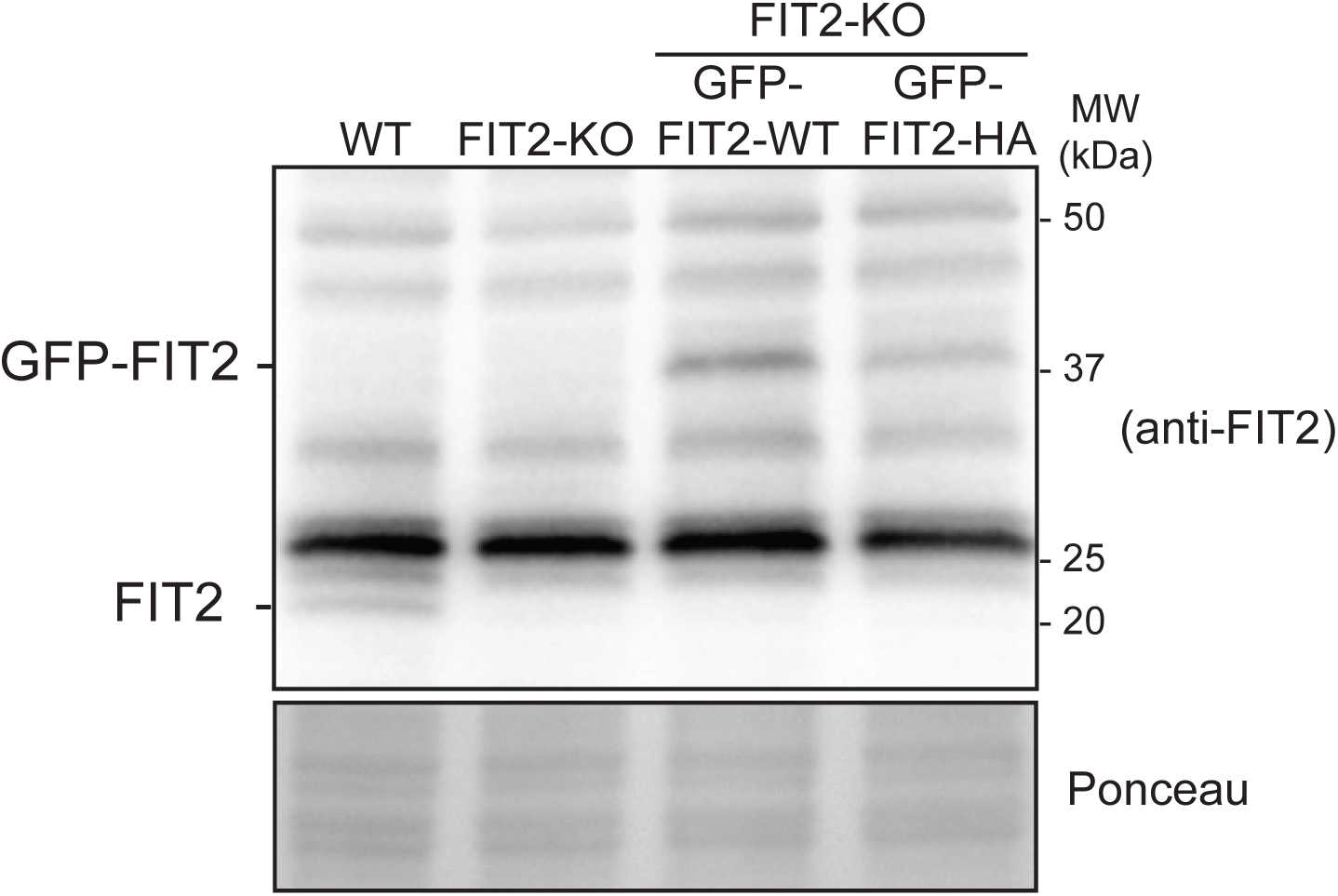
Characterization of FIT2KO SUM159 cells stably expressing GFP-FIT2 wild-type and H214A. Western blot using anti-FIT2 antibody on crude extracts from wild-type cells, FIT2-KO cells, FIT2-KO cells expressing GFP-FIT2 wild-type or H214A under PGK promoter stably integrated in the safe harbor AAVS1 genomic locus. Endogenous FIT2 was observed at the expected size (22 KDa) only in the wild-type cell extract. GFP-tagged forms of FIT2 were observed at the expected size (45 KDa) only in wild-type GFP-FIT2 and H1214A stable cell lines. Ponceau staining was used as loading control.

**Table S1.** Plasmids used in this paper

| Plasmid | Description | Origin/Reference |
| --- | --- | --- |
| pMB98 | pRS416 (CEN,URA3) pADH-GFP (for Nter tagging) | S. Leon lab |
| pMB99 | pRS416 (CEN,URA3) pADH-GFP (for Cter tagging) | S. Leon lab |
| pMB86 | pRS416-derived (CEN, URA3) pADH:Scs3-GFP | This paper |
| pMB87 | pRS416-derived (CEN, URA3) pADH:GFP-Scs3 | This paper |
| pMB96 | pRS416-derived (CEN, URA3) pADH:Scs3-H235A-GFP | This paper |
| pMB97 | pRS416-derived (CEN, URA3) pADH:Scs3-H350A-GFP | This paper |
| pMB104 | pRS416-derived (CEN, URA3) pADH:FIT2-GFP | This paper |
| pMB111 | pRS416-derived (CEN, URA3) pADH:GFP-FIT2 | This paper |
| pMB112 | pRS416-derived (CEN, URA3) pADH:GFP-FIT2-H155A | This paper |
| pMB113 | pRS416-derived (CEN, URA3) pADH:GFP-H214A | This paper |
| pMB193 | pRS426-derived (2μm, URA3) pGAL:FIT2-StrepII | This paper |
| pMB194 | pRS426-derived (2μm, URA3) pGAL:FIT2-H214A-StrepII | This paper |
| pMB65 | pAAVS1-PGK-GFP-FIT2 | This paper |
| pMB68 | pAAVS1-PGK-GFP-FIT2-H214A | This paper |
| pMB69 | pAAVS1-PGK-GFP-FIT2-SGHAAA | This paper |
| pTCW185 | ssRFP-HDEL | F&W lab |
| pJH001 | ssBFP2-KDEL | F&W lab |
| pJH006 | GFP-Sec61 | F&W lab |
|  | LentiCRIPSR-V2 | Addgene #52961 |
|  | psPAX2 | Addgene #12260 |
|  | pCMV-VSVG | Addgene #8454 |
|  | hCas9 | Addgene #41815 |
|  | gRNA-AAVS1-T2 | Addgene #41818 |
|  | ER-ox-GFP | Addgene #68069 |
|  | Emerald-Sec61 | Addgene #54249 |

**Table S2.**
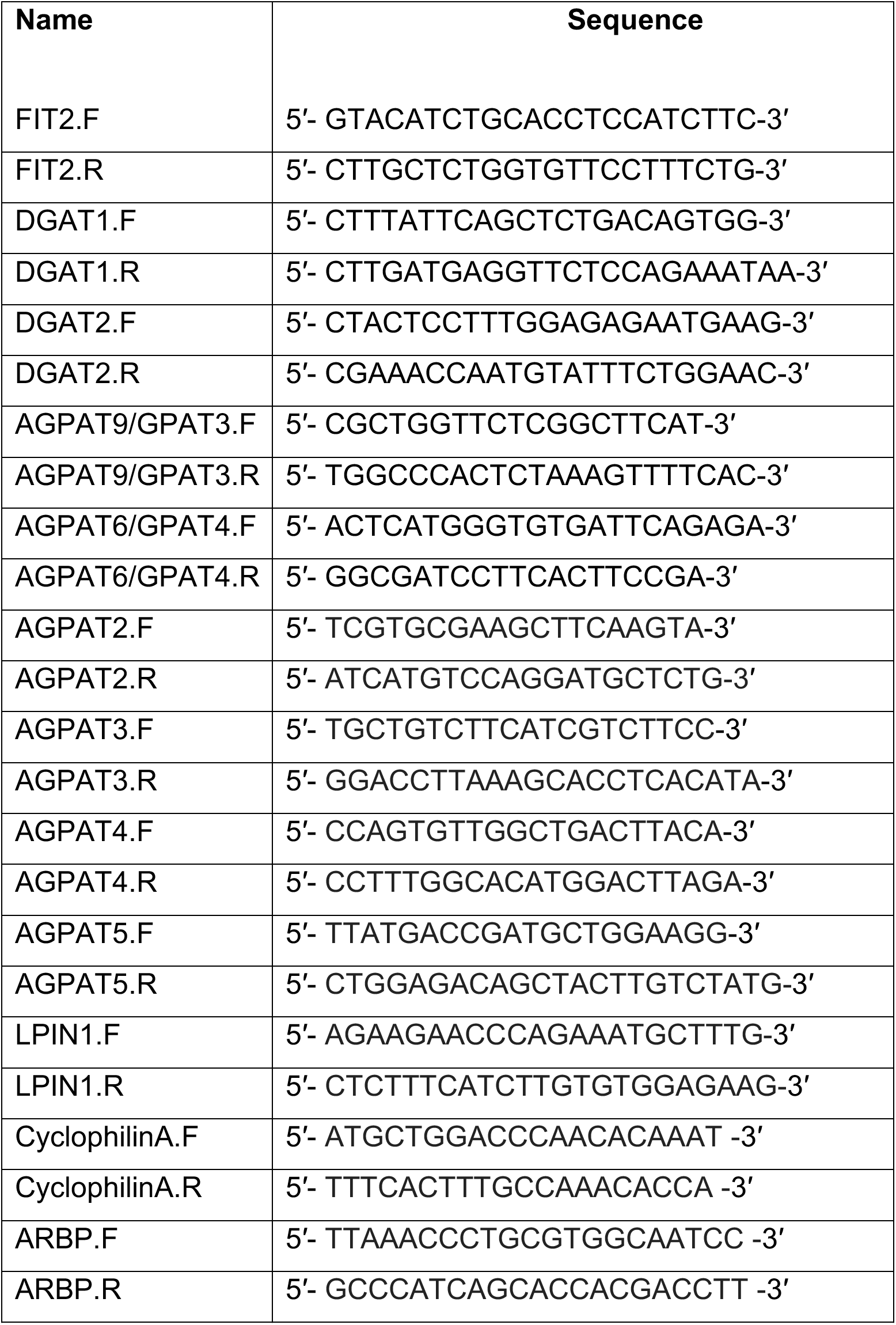
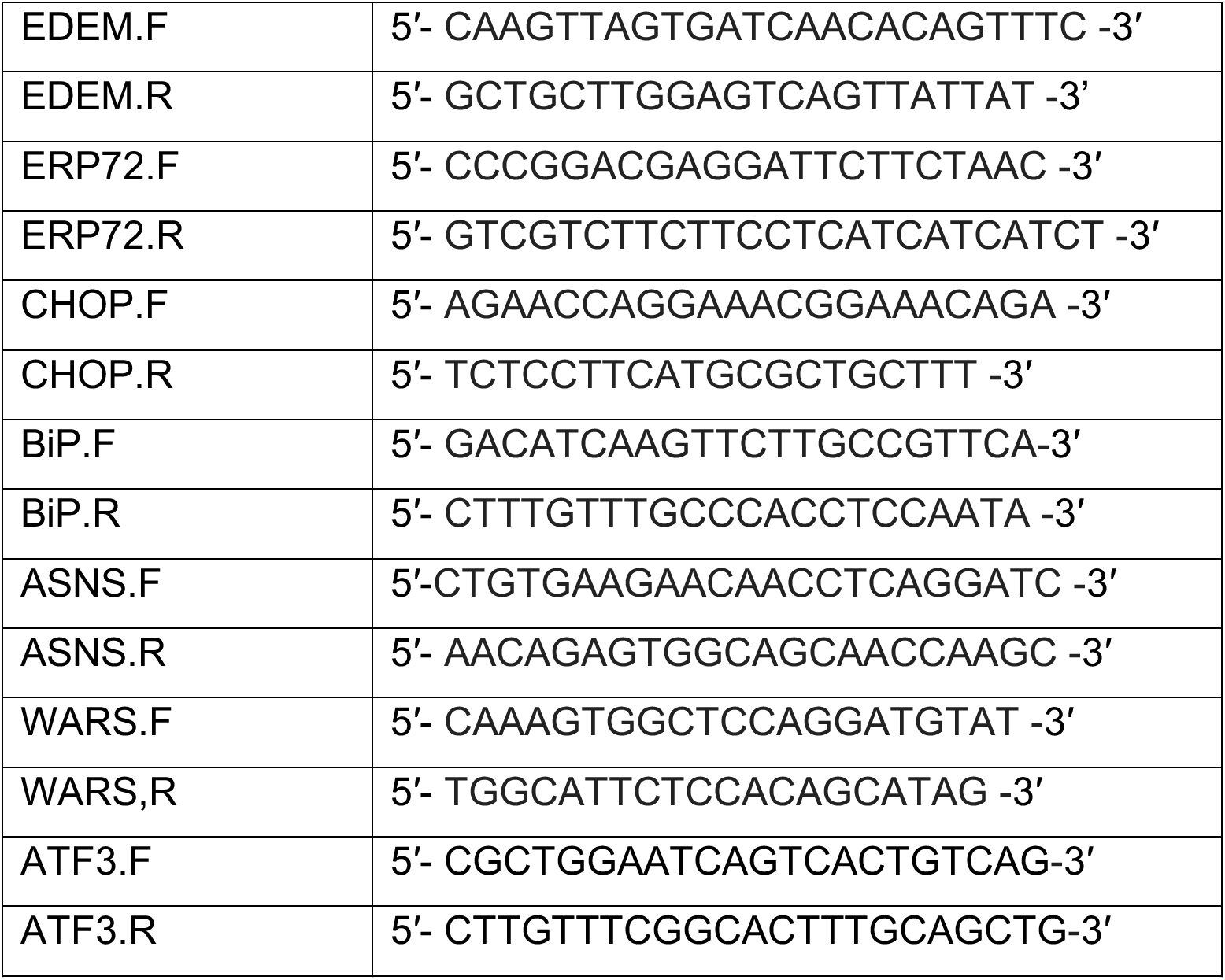
qPCR primers

**Table S3.** Yeast strains

| **Name** | **Genotype** | **Origin/Reference** |
| --- | --- | --- |
| Wild-type<br>(BY4741) | Mat a ; his3Δ1 leu2Δ0 met15Δ0 ura3Δ0 | FW lab |
| scs3Δ | Mat a ; scs3Δ::HisMX leu2Δ0 met15Δ0 ura3Δ0 | This paper |
| iLD | Mat a ; his3Δ0, leu2Δ0, ura3Δ0, lys2Δ0, GalP::DGA1;<br>GalP::ARE2; lro1Δ::HPH®, are1Δ::NAT® | This paper |
| iLD scs3Δ | scs3Δ::HisMX, leu2Δ0, ura3Δ0, lys2Δ0, GalP::DGA1;<br>GalP::ARE2; lro1Δ::HPH®, are1Δ::NAT® | This paper |

## Material and Methods

### Mammalian cell culture, transfection

SUM159 cells (breast carcinoma cell line) were maintained in DMEM/F-12 GlutaMAX (Life Technologies) containing 5% FBS, 100 unit/ml penicillin, and 100 mg/ml streptomycin, 1 mg/ml hydrocortisone (Sigma), 5 mg/ml insulin (Cell Applications, San Diego, CA) and 10 mM HEPES, pH 7.0. Transfection of plasmids was performed 24 h before experiment with Fugene-HD (Promega), according to manufacturer’s instructions. siRNA treatment was performed 72 h before experiment by reverse transfection using Lipofectamine^®^ RNAiMAX (ThermoFisher Scientific), according to manufacturer’s instructions.

### Yeast growth conditions

All yeast strains are derivatives of the BY4741 strain. Yeast was transformed by standard lithium acetate/polyethylene glycol procedure. Cells were grown in synthetic complete (SC) medium containing 2% (wt/vol) glucose, without uracil or leucine to keep plasmid selection. For imaging, cells transformed with the indicated plasmids were grown in SC-URA-LEU medium overnight and imaged during early-log phase (A600=0.3) the next day. For LD induction, iLD strains were precultured to early exponential phase (A600: 0.3) in SC medium containing 2% (wt/vol) glucose and then diluted (A600: 0.05) in SC complete medium containing 2% raffinose (wt/vol) overnight. When cultures reach A600: 0.4–0.6, 2% galactose (wt/vol) was added to the raffinose-grown cells to induce LDs.

### Yeast drop test assay

Wild-type or scs3Δ yeast transformed with the indicated plasmids were grown overnight in SC without uracil liquid culture to stationary phase and spotted in 10-fold serial dilutions onto synthetic media plates without uracil ± inositol ± choline (2 µg/µl). The images shown were taken after 4–5 days of growth at 37ºC.

### Antibodies and Western blots

Rabbit polyclonal antibody against human FIT2 was custom generated (GenScript, Piscataway, NJ), affinity purified and used at a 1:1000 dilution for Western blots. The peptide sequence used for the epitope was the last 16 amino acids “PQSCSLNLKQDSYKKP”. We used antibody against tubulin (T5168, monoclonal, Sigma-Aldrich), calnexin (C5C9, Cell Signaling Technology), eIF2α (9722S; Cell Signaling Technology), Phospho-eIF2α (9721S; Cell Signaling Technology), Ire1α (3294; Cell Signaling Technology), BiP (3177; Cell Signaling Technology). Crude extract samples for Western blots were prepared as described in (Wang et al., 2016). Briefly, cells were lysed with lysis buffer (150 mM NaCl, 50 mM Tris-HCl, pH 7.4, 1% Triton X-100), and spun for 6 min at 16000g. Supernatant was denatured in Laemmli buffer at 37ºC for 10 min. Proteins were separated on 4–15% SDS-PAGE gel (Biorad) and transferred to a PVDF membrane (BioRad).

### RNA extraction and real time-qPCR

Total RNA was isolated using the RNeasy Kit (QIAGEN), according to the manufacturer’s instructions. Complementary DNA was synthesized using iScript cDNA Synthesis Kit (Bio-Rad), and qPCR was performed in triplicates using Power SYBR Green PCR Master Mix Kit (Applied Biosystems). Sequences of the qPCR primers used are listed in Table S2.

### Plasmid constructs

Plasmids in this paper are described in Table S1. To obtain pADH:Scs3-GFP (pMB86) and pADH:GFP-Scs3 (pMB87), *SCS3* gene was amplified from BY4741 genomic DNA, inserted into pCRII-ZeroBlunt (Invitrogen) (pMB88), and subcloned into XbaI–EcoRI sites in pRS416 (pMB98) to make pMB86 and into XhoI-EcoRI in pRS416 (pMB99) to make pMB87. Site-directed mutagenesis (QuikChange Lightning Site-Directed Mutagenesis Kit, Agilent) was performed by PCR using mutagenic oligonucleotides with pMB88 as template. Mutagenized plasmids were checked by sequencing, and a region containing the mutation was systematically subcloned into the original pMB98 using endogenous restriction sites to prevent mutations that may have occurred elsewhere on the plasmid. Mutation on pMB86 of the C2 and C3 conserved histidine to alanine led to pMB96 and pMB97, respectively. The pADH:FIT2-GFP (pMB104) and pADH:GFP-FIT2 (pMB111) were amplified from human FITM2 cDNA NM_001080472.2 (Origene), and subcloned into XbaI–EcoRI sites in pRS416 (pMB98) to make pMB104 and into XhoI-EcoRI sites in pRS416 (pMB99) to make pMB111. Mutation on pMB111 of the C2 and C3 conserved histidine to alanine led to pMB112 and pMB113, respectively. Plasmids used for protein purification were derived from pRS426 pGAL:TAP. To obtain pGal:FIT2-StrepII, FIT2 gene from pMB104 was first cloned into SpeI/EcoRI sites. StrepII tag (WSHPQFEK), obtained from a synthetic fragment (gBlock, Addgene), was then cloned into EcoRI/XhoI. Same strategy was used to obtain pGal:FIT2-H214A-StrepII, using pMB112 as template for FIT2-H214A gene. Plasmids used for safe harbor integration were derived from AAVS1-PGK1 HR donor plasmid (Addgene #68375). Wild-type GFP-FIT2 and H155A sequences were cloned using XbaI/KpnI restriction sites. Same plasmids were also used for transient transfection in SUM159 cells.

### Generating cell line knock-out and knock-in in SUM159 cells

To generate CRISPR FIT2-KO SUM159 cell line, we used lentiviral delivery method as described in (Sanjana et al., 2014). The sequence 5′-CGGGGTGCACTCACACGTTG-3′ was used as a gRNA to direct Cas9 into the exon 1 of the FIT2 locus. gRNA was cloned in LentiCRISPR V2 plasmid (Addgene #52961). Briefly, LentiCRISPR V2 plasmid, psPAX2 (Addgene #12260) and pCMV-VSVG (Addgene #8454) plasmids were transfected in HEK293 packaging cells and lentiviral particles were collected from culture media 24h later and used to infect SUM159 cells. 1 day after infection, cells were treated with puromycin for 3 days to select for positively infected cells. Cells were re-plated into 150 mm dishes at clonal density. Individual colonies were then isolated in 24-well dishes. Screening of positive clones was performed by qPCR (sense primer: 5′-GTACATCTGCACCTCCATCTTC-3′ and antisense primer 5′-CTTGCTCTGGTGTTCCTTTCTG-3′) using power SYBR green (Life Technologies). Genomic DNA of clones showing mRNA expression defect were extracted (Quick Extract DNA extraction solution; Epicentre), and the genomic sequence surrounding the target exon of FIT2 was amplified by PCR (sense 5′-TCAAGCTGTGGGACTCGCGG-3′ and antisense 5′-TCCAATGACTCGTCCACCAC-3′). PCR products were subcloned into a plasmid (Zero Blunt TOPO PCR cloning kit; Life Technologies) to validate the edited region of positive knockout clones by sequencing. Knock-out was also confirmed by western blot using costumed FIT2 antibody.

To generate cell line stably expressing GFP-FIT2 in FIT2-KO SUM159 cells, we used AAVS1 Safe Harbor targeting method (System Biosciences). Wild-type GFP-FIT2 or H155A donor construct was co-transfected with hCas9 plasmid and gRNA-AAVS1-T2 plasmid using Lipofectamine 3000 (ThermoFisher Scientific), according to manufacturer’s instructions. Cells were selected with puromycin over 3 days, and single-cell FACS sorting was performed, based on GFP signal. Positive clones were confirmed by fluorescent microscopy and by western blot with anti-FIT2 antibody.

### Protein purification

Purification protocol was adapted from (Gross and Silver, 2013). The *S. cerevisiae* strain BY4741 was used for the galactose-inducible expression of StrepII tagged human FIT2 protein. For galactose induction, 600 ml of yeast cells were precultured to early exponential phase (A600: 0.3) in SC-URA medium, containing 2% raffinose (wild-type/vol) and 0.02% glucose (wild-type/vol) to initiate growth. The culture was then diluted into SC-URA medium (6 liters) with 2% galactose as the carbon source to induce the expression of StrepII-tag FIT2. Cells were harvested after overnight growth (A600: approx. 2), and pellet was stored in -80° C. Cells were disrupted with LM20 Microfluidizer^®^ High Shear Fluid Processor (4 runs, 30000 psi) in lysis buffer containing 50 mM Tris-HCl, pH 7.5, 200 mM NaCl, 10% glycerol and supplemented with Complete protease inhibitor cocktail (Roche). Unbroken cells were removed by centrifugation at 10,000g for 30 min. The total cell lysate (supernatant) was ultracentrifuged at 100,000g for 90 min. Membrane fraction (pellet) was resuspended in lysis buffer supplemented with 1% Fos-Choline 13 and incubated overnight at 4° C, conditions described to solubilize FIT2 (Gross and Silver, 2013). Unsolubilized membranes were removed by ultracentrifugation at 100,000g for 60 min. Supernatant fraction containing solubilized FIT2 was incubated with Strep-Tactin superflow plus beads (Qiagen) for 3 hours. Sample was then transferred to 25-ml BioSpin columns and washed by gravity flow with 150–200 ml of lysis buffer supplemented with 0.1% Fos-Choline 13. Protein was eluted in 5 min sequential elutions in 500 μl of lysis buffer containing 0.1% Fos-Choline 13 and 10 mM desthiobiotin (Sigma). Eluted fractions were pooled and concentrated using 10-kDa Amicon Ultra-4 Centrifugal Filter Units (Millipore). Protein concentration was calculated by comparing band intensity of 2µl of eluate with BSA standard on 4–15% SDS-PAGE gels (Biorad) and visualized by Coomassie blue (Imperial Protein Stain, ThermoFisher Scientific). Aliquots were stored at 80° C.

### Enzymatic assays

Lysophosphatidic acid phosphatase (LPA) activity was measured by following the formation of monoacylglycerol (MAG) from soluble lysophosphatidic acid, L-a-oleoyl [oleoyl-1-^14^C] (American Radiolabeled Chemicals). For time-dependent reactions, the reaction mixture contained 20 nmoles of cold C18:1n9 LPA (Avanti Polar lipid) and 2 nmoles radioactive LPA, dried under airflow and sonicate in reaction buffer 50 mM Tris-HCl (pH 7.5), 50 mM NaCl, 0.5 mM MgCl_2_, 0.02% Triton X-100. 100 ng of enzyme protein was added in a total volume of 0.1-ml reaction mixture and incubated at 37° C for the indicated time. For substrate dependent reactions, series of twofold dilution were performed in reaction buffer and incubated with 100 ng of wild-type FIT2 or H214A proteins, or Mock eluate for 1 h at 37ºC. Reaction was stopped by adding chloroform:methanol mixture (2:1) and phosphoric acid 2%. After centrifugation 5 min at 5000g, lower phase was extracted, dried under air stream, resuspended in chloroform and loaded on TLC using chloroform:methanol (24:1) solvent system. TLC plate were exposed to phosphor imaging cassette overnight and revealed with Typhoon FLA 7000 phosphor imager. MAG bands were scraped and quantified by liquid scintillation counter. Enzyme assays were conducted in triplicate, and the average values and S.D. of the assays were corrected against Mock extract control samples. Similar assays were performed to determine phosphatidic acid (PA) phosphatase activity; the formation of diacylglycerol (DAG) from mixture containing 20 nmoles of cold egg PA (Avanti Polar lipid) and phosphatidic acid, L-a-dioleoyl [oleoyl-1-^14^C] (American Radiolabeled Chemicals) was measured via TLC and scintillation counting. For lysophosphatidylcholine lipase assay, 25 nmoles of 1-oleoyl-sn-glycero-3-phosphocholine (Sigma) was mixed with 2 nmoles of lysophosphatidylcholine, L-a-oleoyl [oleoyl-1-^14^C] (American Radiolabeled Chemicals) in 100 μl of reaction mixture, and incubated with 100 ng of wild-type FIT2 or H214A proteins, or Mock eluate for 1 h at 37ºC. Indicated units of phospholipase C from *Clostridium perfringens* (Sigma), resuspended in reaction buffer, were used as positive controls.

### [^14^C]-Oleic acid labeling of lipids, lipid extraction and TLC

Cells were pulse-labeled with [^14^C]-oleic acid (50 µCi/µmol), washed with phosphate buffer saline three times. Lipids were extracted directly from six-well cell-culture plates by adding hexane:isopropanol mixture (3:2) and gentle shaking for 10 min. The process was repeated a second time for full extraction of all lipids. Lipids were dried under nitrogen stream and separated by TLC using hexane:diethyl ether:acetic acid (80:20:1) solvent system for neutral lipid separation, and using chloroform:methanol:acetone:acetic acid:water (50:10:20:15:5) solvent system for phospholipid separation. TLC plates were exposed to phosphor imaging cassette overnight and revealed by Typhoon FLA 7000 phosphor imager. Incorporation of radiolabeled oleic acid in lipids was determined over time by measuring band intensity on TLC using Image J software.

### Lipidomics

SUM159 cells were grown on a 10-cm culture dish treated or not with 500 μM oleic acid for 24h until they reached ~80% confluency. Subsequently, lipids were extracted according to Folch’s Method (Folch et al., 1957). The organic phase of each sample, normalized by total protein (using BCA assay), were then separated by an HPLC-MS method adopted from (Bligh and Dyer, 1959). Briefly, HPLC analysis was performed employing a C30 reverse-phase column (Thermo Acclaim C30, 2.1 × 250 mm, 3 μm, operated at 55° C; Thermo Fisher Scientific) connected to a Dionex UltiMate 3000 HPLC system and a QExactive orbitrap mass spectrometer (Thermo Fisher Scientific) equipped with a heated electrospray ionization (HESI) probe. Dried lipid samples were dissolved in corresponding volumes of 2:1 methanol:chloroform (v/v) and 5 μl of each sample was injected, with separate injections for positive and negative ionization modes. Mobile phase A consisted of 60:40 water/acetonitrile, including 10 mM ammonium formate and 0.1% formic acid, and mobile phase B consisted of 90:10 2-propanol/acetonitrile, also including 10 mM ammonium formate and 0.1% formic acid. The elution was performed with a gradient of 90 minutes; during 0–7 minutes, elution starts with 40% B and increases to 55%; from 7 to 8 minutes, increase to 65% B; from 8 to 12 minutes, elution is maintained with 65% B; from 12 to 30 minutes, increase to 70% B; from 30 to 31 minutes, increase to 88% B; from 31 to 51 minutes, increase to 95% B; from 51 to 53 minutes, increase to 100% B; during 53 to 73 minutes, 100% B is maintained; from 73 to 73.1 minutes, solvent B was decreased to 40% and then maintained for another 16.9 minutes for column re-equilibration. The flow-rate was set to 0.2 mL/min. The column oven temperature was set to 55° C, and the temperature of the autosampler tray was set to 4° C. The spray voltage was set to 4.2 kV, and the heated capillary and the HESI were held at 320° C and 300° C, respectively. The S-lens RF level was set to 50, and the sheath and auxiliary gas were set to 35 and 3 units, respectively. These conditions were held constant for both positive and negative ionization mode acquisitions. External mass calibration was performed using the standard calibration mixture every 7 days. MS spectra of lipids were acquired in full-scan/data-dependent MS2 mode. For the full-scan acquisition, the resolution was set to 70,000, the AGC target was 1e6, the maximum integration time was 50 msec, and the scan range was m/z = 133.4–2000. For data-dependent MS2, the top 10 ions in each full scan were isolated with a 1.0 Da window, fragmented at a stepped normalized collision energy of 15, 25, and 35 units, and analyzed at a resolution of 17,500 with an AGC target of 2e5 and a maximum integration time of 100 msec. The underfill ratio was set to 0. The selection of the top 10 ions was subject to isotopic exclusion with a dynamic exclusion window of 5.0 sec. Processing of raw data was performed using LipidSearch software (Thermo Fisher Scientific/Mitsui Knowledge Industries) (Taguchi and Ishikawa, 2010; Yamada et al., 2013). The assembled results were exported to R-Studio where all identified lipids were included for subsequent analyses if they fulfilled the following LipidSearch-based criteria: 1) reject equal to zero, 2) main grade A OR main grade B AND APValue<0.01 for at least three replicates, and 3) no missing values across all samples. Based on these filters, 494 out of 990 lipids (without oleate treatment) and 541 out of 987 lipids (with oleate treatment) were present in all samples and displayed high accuracy of identification. Further quality controls were performed using pair-wise correlations between replicates and principal component analysis (PCA, FactomineR package (Le et al., 2008), comparing sample groups. All replicates displayed high reproducibility (r-value>0.77 and p-value<0.01) and the PCA distinguished between sample groups on the first and second principal components with 95% confidence interval (results not shown).

To enrich for phosphatidates, we followed the extraction and separation protocol developed in (Triebl et al., 2014). Briefly, cells (one 10-cm dish, 80-90% confluent was used for each replicate) treated or not with 500 μM oleic acid for 24h, were resuspended and lysed by sonication/snap freezing cycle in 0.1M HCl. The lipid (lower) phase of each sample, normalized by total protein (using BCA assay), was extracted with 0.1M HCl:Methanol 1:2(v/v), vortex for 1 min, then 1 volume of chroloform was added, vortex for 1min and centrifuge at 1000g for 5 min. Lower phase was dried under nitrogen flow and re-suspended in 70 µL chloroform:methanol 1:1 (v/v). To separate phosphatidates, HPLC-MS analysis was performed employing a Kinetex HILIC column (2.1 mm × 100 mm, 2.6 m, operated at 55˚ C; Phenomenex), connected to a Dionex UltiMate 3000 HPLC system and a QExactive orbitrap mass spectrometer (Thermo Fisher Scientific) equipped with a heated electrospray ionization (HESI) probe. Mobile phase A consisted of deionized water containing 10 mM ammonium formate and 0.5% formic acid. Mobile phase B consisted of 2-propanol/acetonitrile 5:2 (v/v) containing 10 mM ammonium formate and 0.5% formic acid as well. Gradient elution started at 5% A with a linear increase to 50% A over the course of 12 min followed by an isocratic elution with 50% A for 3 more min. Finally, the column was re-equilibrated for 15 min. The flow rate was set to 0.3 mL/min. Sample solvent was chloroform/methanol 1:1 (v/v) and 20 µL were injected. The temperature for the autosampler tray was set to 4° C. The spray voltage was set to 3.5 kV, and the heated capillary and the HESI were held at 350° C and 300° C, respectively. The S-lens RF level was set to 50, and the sheath and auxiliary gas were set to 20 and 5 units, respectively. Data were acquired in negative ionization mode. The Orbitrap mass spectrometer was operated in data dependent analysis mode; a full scan spectrum was obtained in the Orbitrap mass analyzer at R = 70,000 while simultaneously tandem mass spectra of the ten most abundant full scan masses (selected from an inclusion list containing phosphatidic acid and lysophosphatidic acid molecular species) were obtained in the linear ion trap at a normalized collision energy of 35% and an exclusion duration of 10s.

### Live cell imaging and image processing

Microscopy was performed on spinning disk confocal (Yokogawa CSU-X1) set up on a Nikon Eclipse Ti inverted microscope with a 100 ApoTIRF 1.4 NA objective (Nikon, Melville, NY) in line with 2x amplification. Fluorophores were excited with 405-, 488-, or 561-nm laser lines, and fluorescence was detected by an iXon Ultra 897 EMCCD camera (Andor, Belfast, UK) or Zyla 4.2 Plus sCMOS camera (Andor, Belfast, UK). Bandpass filters (Chroma Technology) were applied to all acquisitions. Where applicable, Z stacks of 0.15-mm slices were obtained with piezo Z-stage. For live-cell imaging, temperature, humidity and CO_2_ were controlled during imaging using a stage-top chamber (Oko Lab). For imaging of initial LD formation or late events, 500 µM oleic acid (Sigma) complexed with 0.5% fatty acid-free BSA (Sigma) were added to wells immediately before image acquisition. Where applicable, 0.5 mg/ml BODIPY 493/503 (Life Technologies) or HCS LipidTOX™ Deep Red Neutral Lipid Stain (ThermoFisher Scientific) was added before and with oleic acid supplementation to stain LDs. For DGATs inhibitor experiments, cells were treated 2 h before oleate addtion with 10 µM inhibitor. Inhibitors were obtained from collaborators at Merck & Co., Inc. and already published elsewhere (Imbriglio et al., 2015; Liu et al., 2013).

Acquired images were processed and quantified manually with FIJI software (http://fiji.sc/Fiji). The plugin “Find Maxima” was used to quantify yeast LD number. The software Cell Profiler was used to quantify LD number in SUM159 cells experiments using automated pipeline. All quantifications were done on raw images. Where necessary, deconvolution of images was performed using Huygens Professional 15.05 (Scientific Volume Imaging) with CMLE algorithm using a calculated PSF for each wavelength.

### Quantification of endoplasmic reticulum sheets and tubules

0.25 μm Z-stack images were acquired on confocal spinning-disk from cells expressing the ER marker oxGFP-KDEL. From these images, sheets and tubules were quantified from masks obtained via supervised segmentation. The pipeline has the following steps: 1. A few volume crops are annotated, with a subset of their voxels being labeled as background, tubules, or sheets. 2. A machine learning (random forest) model (available here https://github.com/HMS-IDAC/VoxelClassifier) is trained to classify voxels based on provided annotations. 3. A dataset of crops to be analyzed is generated -- since voxel classification in 3D is computationally time-consuming, this step speeds up analysis considerably. 4. The trained machine learning model is applied on crops from step 3 to generate masks with segmented sheet/tubule areas. 5. An algorithm based on Watershed (Roesdink and Meijster, 2001) and Active Contours (Chan and Vese, 2001) is deployed on the crops and the corresponding probability maps to create nuclei and cell masks. 6. Layer masks of thickness approximately 25 voxels are generated, outwardly from the nucleus, using morphological dilation, and the number of sheets and tubules in such masks is computed, up to 20 layers, or until the amount of sheet voxels stops growing, whichever happens first. 7. Step 6 is repeated after the crops and probability maps are re-sized so that the cell mask volume is equal to the average cell volume in the analyzed dataset.

### Immunofluorescence

For standard immunofluorescence experiments, cells were grown on glass bottom 6-well plates and fixed with 4% formaldehyde. Blocking was performed with 3% BSA in PBS + 0.1% Triton X-100. Primary and secondary antibody dilutions were performed in the same solution. We used anti-LAMP1(D2D11; Cell signaling Technology) to detect lysosomes, anti-catalase (D4P7B; Cell Signaling Technology) to detect peroxisomes, anti-GM130 (D6B1; Cell Signaling Technology) to detect Golgi apparatus. We used as a secondary antibody Alexa 568–conjugated secondary antibodies (Invitrogen). Nuclei were stained with 1 μg/ml Hoechst 33342 (Invitrogen) during one of the post-secondary antibody washes.

### Electron microscopy

Cells in petri dishes were fixed in 2.5% gluteraldehyde in 0.1 M sodium cacodylate buffer, pH 7.4, for 1 h. Buffer rinsed cells were scraped in 1% gelatin and spun down in 2% agar. Chilled blocks were trimmed and postfixed in 1% osmium tetroxide for 1 h. The samples were rinsed three times in sodium cacodylate rinse buffer and postfixed in 1% osmium tetroxide for 1 h. Samples were then rinsed and en bloc stained in aqueous 2% uranyl acetate for 1 h, followed by rinsing, dehydrating in an ethanol series and infiltrated with Embed 812 (Electron Microscopy Sciences) resin and baked over night at 60⁰C. Hardened blocked were cut using a Leica UltraCut UC7. Sections (60 nm) were collected on formvar/carbon-coated nickel grids and contrast stained with 2% uranyl acetate and lead citrate. They were viewed using an FEI Tencai Biotwin TEM at 80Kv. Images were taken on a Morada CCD using iTEM (Olympus) software.

### Lattice Light-Sheet microscopy

Imaging was performed on a lattice light-sheet microscope developed by (Chen et al., 2014) and built by Intelligent Imaging Innovations, Inc (3i). SUM159 cells were transfected with the ER marker ox-GFP (Addgene #68069) and imaged at 37° C in Leibovitz’s L-15 Medium (Thermo Fisher Scientific, 11415064) supplemented with 5% heat-inactivated FBS (GE Healthcare Life Sciences SH30071.03), 100 units/mL penicillin and 100 µg/mL streptomycin (Thermo Fisher Scientific 15140122), 1 µg/mL hydrocortisone (Sigma-Aldrich H0888), 25 mM HEPES (Thermo Fisher Scientific 15630080), and 5 µg/mL insulin (Cell Applications 128). A square lattice light sheet was generated and passed through an annular mask with inner and outer NAs of 0.42 and 0.50, respectively. The beam was dithered through the sample during acquisition. Cell volumes were acquired by stepping the sample stage at 400nm intervals between 221 and 251 times with a 35-ms exposure at each plane. Cells were excited with a 488-nm laser (~300 mW power; ~15.87 µW-28.9 µW measured at the back aperture of the excitation objective during experiments). Fluorescence was detected with a 25x 1.1NA Nikon CFI APO LWD Objective, passed through a 488-nm StopLine single-notch filter (Semrock NF03-488E-25), a 446/523/600/677-nm Brightline quad-band bandpass filter (Semrock FF01-446/523/600/677-25), and a custom dichroic transmitting 390–555 nm, 645–875 nm and reflecting 565–630 nm onto a Hammamatsu ORCA Flash 4.0 sCMOS camera. Images were captured as 1024×256 pixel planes and subsequently deskewed and background corrected using 3i’s SlideBook software. Image data was loaded into Matlab using code developed in Tom Kirchhausen lab at Harvard Medical School (Aguet et al., 2016) and subsequently deconvolved by providing measured background and an experimentally measured PSF for 15 iterations using a Richardson-Lucy algorithm adapted to run on a graphics processing unit developed by Eric Betzig Lab at Janelia.

### Statistical analysis

All results were analyzed using Prism (GraphPad Software), and the tests specified in the figure legends, unless otherwise stated. For figure 2A and 2C, data were exported to R-Studio where 90% confidence ellipses were plotted using ggplot2 (Wickam, 2010). All error bars represent the standard deviation.

